# Intravenous gene therapy improves lifespan and clinical outcomes in feline Sandhoff Disease

**DOI:** 10.1101/2024.11.15.623838

**Authors:** Anne S. Maguire, Linh Ta, Amanda L. Gross, Devin E. Osterhoudt, Jessica S. Cannon, Paige I. Hall, Maninder Sandey, Thomas N. Seyfried, Heather L. Gray-Edwards, Miguel Sena-Esteves, Douglas R. Martin

## Abstract

Sandhoff Disease (SD), a fatal neurodegenerative disorder, is caused by the absence of ß-hexosaminidase (Hex) and subsequent accumulation of GM2 ganglioside in lysosomes. Previous studies have led to adeno-associated virus (AAV) gene therapy for children with GM2 gangliosidosis in both expanded access and Phase I/II clinical trials via intracranial and/or cerebrospinal fluid-based delivery. The current study investigated intravenous (IV) gene therapy of SD cats, treated at one month of age with a bicistronic AAV vector. While untreated SD cats lived to 4.3±0.2 months, cats treated with low and high doses lived to 8.3±1.2 and 12.4±2.7 months, respectively. In-life assessments revealed clear clinical benefit of AAV treatment, with the most dramatic improvement seen in the reduction of overt full-body tremors. Cerebrospinal fluid levels of aspartate aminotransferase (AST) and lactate dehydrogenase (LDH) were decreased, indicating a reduction of cell damage within the central nervous system. Magnetic resonance imaging (MRI) and spectroscopy (MRS) acquired on a 7 Tesla scanner indicated that structural pathology and metabolite abnormalities are partially normalized by AAV treatment. Dose-dependent reduction of GM2 ganglioside storage and increases in Hex activity were most substantial in the caudal regions of the brain and in the spinal cord. Immunohistochemistry revealed reduction in neuroinflammatory cell populations and partial correction of myelin deficits. These results support the dose-dependent efficacy of AAV delivered IV for significant restoration of clinical metrics and Hex function in a feline model of SD.

**One Sentence Summary:** Intravenous administration of AAV gene therapy is safe and efficacious in a feline model of Sandhoff disease.

## INTRODUCTION

Sandhoff Disease (SD) and Tay-Sachs Disease (TSD) are neurodegenerative lysosomal storage diseases (LSDs) that result in the death of affected children by 4 years of age. Because there are no FDA-approved therapies, current treatment strategies are limited to palliation. SD and TSD are clinically similar forms of GM2 gangliosidosis (GM2), and their pathogenesis results from the absence of β-hexosaminidase (Hex) and subsequent accumulation of GM2 ganglioside in lysosomes. The deficiency of Hex activity results in a cascade of pathological effects throughout the body, with the most severe dysfunction in the central nervous system (CNS).

Though enzyme replacement, stem cell, and chaperone therapies (*1*) continue to be thoroughly explored as treatment strategies for SD and TSD, none have achieved the level of efficacy in animal models that gene therapy has. Through injection of adeno-associated viral (AAV) vectors, Hex activity and GM2 ganglioside storage deficits are consistently improved with a variety of capsids, delivery routes, and animal models (*2–7*). AAV1, AAV2 and AAVrh8 were used for direct injections of the brain in early investigations, with treatment consisting of both subunits of Hex (α and β) expressed from separate vectors in a 1:1 mixture (*4, 8*). Bilateral injection of AAVs into the thalamus and deep cerebellar nuclei (DCN) of affected cats quadrupled lifespan and completely abrogated full-body tremors, the most debilitating feature of feline SD. In fact, AAV injection of the CNS was so effective that approximately half of the treated SD cats died of emergent peripheral organ disease, which remains subclinical in untreated cats due to their severe and rapidly progressive CNS dysfunction (*9*).

Concerns about injection into the highly vascular posterior fossa, where the cerebellum resides, led to the investigation of less invasive routes. Several reports suggest that cerebrospinal fluid (CSF) is an effective substitute for DCN injection in mice, cats and sheep (*7, 10, 11*). AAV delivery via CSF was refined in sheep by using a flexible catheter inserted at the lumbar spinal cord and advanced rostrally into the cisterna magna (*12*). The studies described above led to expanded access (*13*) or Phase I/II clinical trials in children with GM2 gangliosidosis using AAV vectors expressing the Hex α and β subunits delivered to the thalamus and CSF or to the CSF alone (ClinicalTrials.gov Identifiers NCT04669535, NCT04798235).

Intravenous (IV) delivery of gene therapy is a less invasive approach that may effectively treat the CNS when using a capsid, such as AAV9, that can penetrate the blood brain barrier (BBB) (*14*). Though debate continues about the pros and cons of IV versus CSF-based treatment approaches for CNS disorders (*15*), numerous reports in animal models demonstrate the efficacy of AAV delivered by the IV route (*6, 16–21*), and IV-delivered AAV9 is approved by the U.S. Food and Drug Administration (FDA) and the European Medicines Agency (EMA) for treatment of spinal muscular atrophy (*22*). Positive results of a phase I/II clinical trial of AAV9 delivered IV to children with GM1 gangliosidosis (NCT03952637) further support the potential of this approach. However, simultaneous expression of both the α and ß subunits of Hex is essential for optimal production of HexA (*23–25*), the isozyme responsible for GM2 ganglioside degradation in humans. Delivery of 2 separate vectors expressing each Hex subunit is not a viable strategy for IV injection due to vector dilution in the bloodstream and low likelihood that cells will be transduced by both vectors. Thus, the purpose of the current study was to test a bicistronic vector expressing both Hex subunits (*19*) for scale-up in the feline model of SD and potential translation to human patients. If effective, the bicistronic vector expressing both Hex subunits could be used to treat both Sandhoff disease and Tay-Sachs disease.

## RESULTS

### Improved lifespan and quality of life

SD cats were treated by IV injection of a bicistronic AAV9 vector expressing both the α and β subunits of Hex as previously described (*19*), except that feline-specific cDNAs were used (see Materials and Methods). A single injection of vector at 5×10^13^ vg/kg (low dose) or 2×10^14^ vg/kg (high dose) was performed through a peripheral vein over approximately 2 minutes. No sedation was required.

The lifespan of SD cats was doubled or tripled by AAV treatment with the low or high dose, respectively (untreated: 4.3±0.2 months, low dose: 8.3±1.2 months, high dose: 12.4±2.7 months). The longest-lived cat (ID: 55) was 16.0 months old at humane endpoint, defined as the point at which the cat can no longer stand, usually due to pelvic limb weakness / paresis (Fig. 1A). Moreover, AAV treatment improved the quality of life in SD cats as measured by weight (Fig. 1B) and a neurological clinical rating score (Fig. 1C) in a dose-dependent fashion, though even the high dose failed to normalize both metrics. The most dramatic clinical improvement was the reduction of body tremors, the most debilitating feature of feline SD. While untreated SD cats developed overt full-body tremors by 2.4±0.1 months, SD cats treated with the high dose never developed full-body tremors. In the low-dose cohort, 1 of 4 cats did not develop full-body tremors while the remaining 3 cats had a delayed onset at 6.8±1.8 months of age (p=0.034) (Table 1). Cats in the high dose long-term cohort reached humane endpoint through pelvic limb paresis (Table 1) due to spinal cord compression (Fig S1), instead of debilitating full-body tremors as in untreated SD cats.

**Figure 1.**
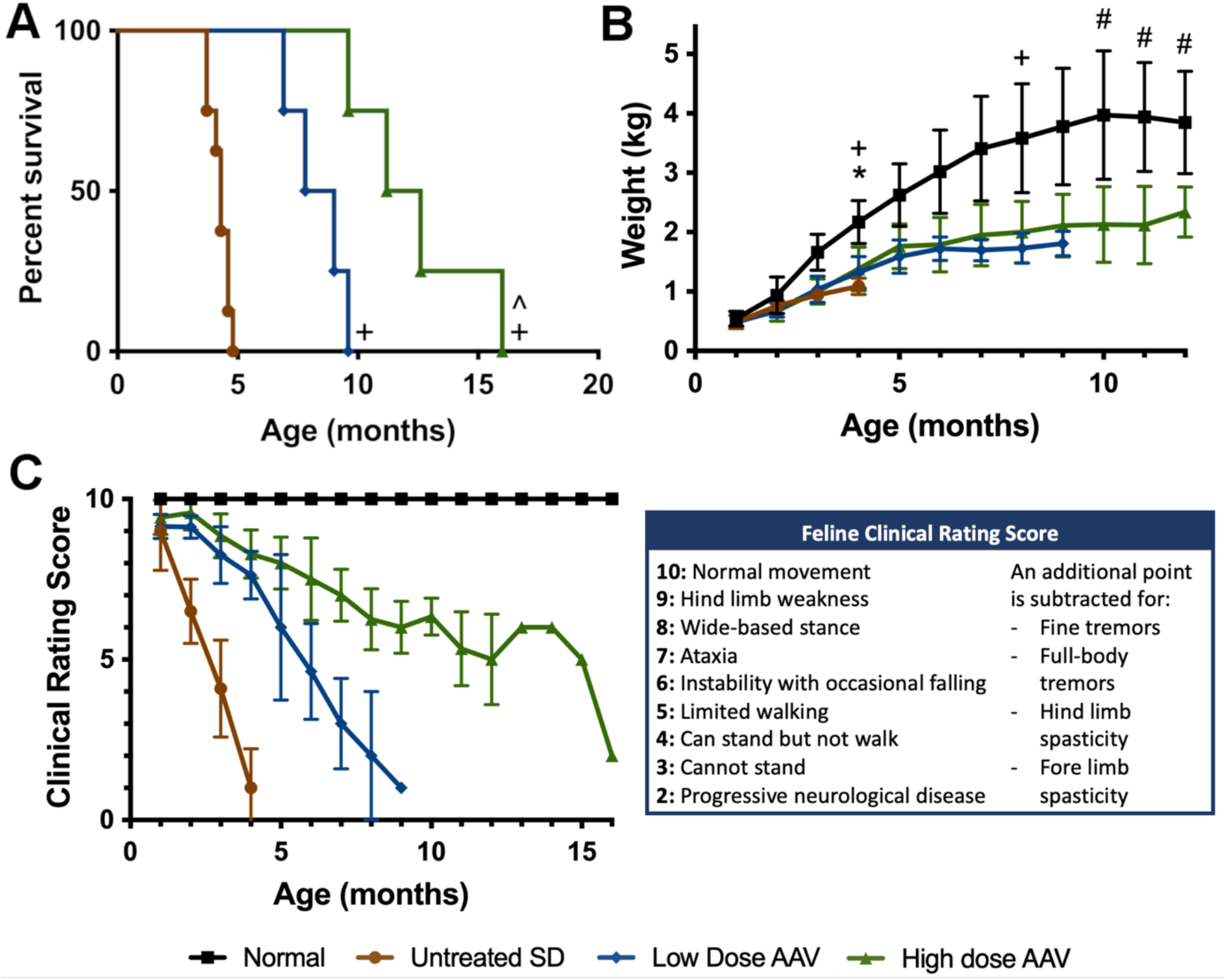
Survival and quality of life improve with AAV treatment. (A) Kaplan-Meier curve demonstrates improved survival with AAV treatment. The lifespan of cats treated with the low dose and high dose nearly doubled and tripled, respectively. ^+^p<0.05 v. untreated SD, ^p<0.05 v. low dose. (B) Weights of treated cats were partially normalized. *p<0.05 untreated SD v. normal; ^+^p<0.05 low dose v. normal; ^#^p<0.05 high dose v. normal. (C) Neurological clinical rating scores (CRS) indicate dose-dependent delay of neurologic dysfunction. CRS for the high-dose cohort (green) artifactually increased at 13 months because only one cat with better-than-average treatment survived to this age.

**Table 1.**
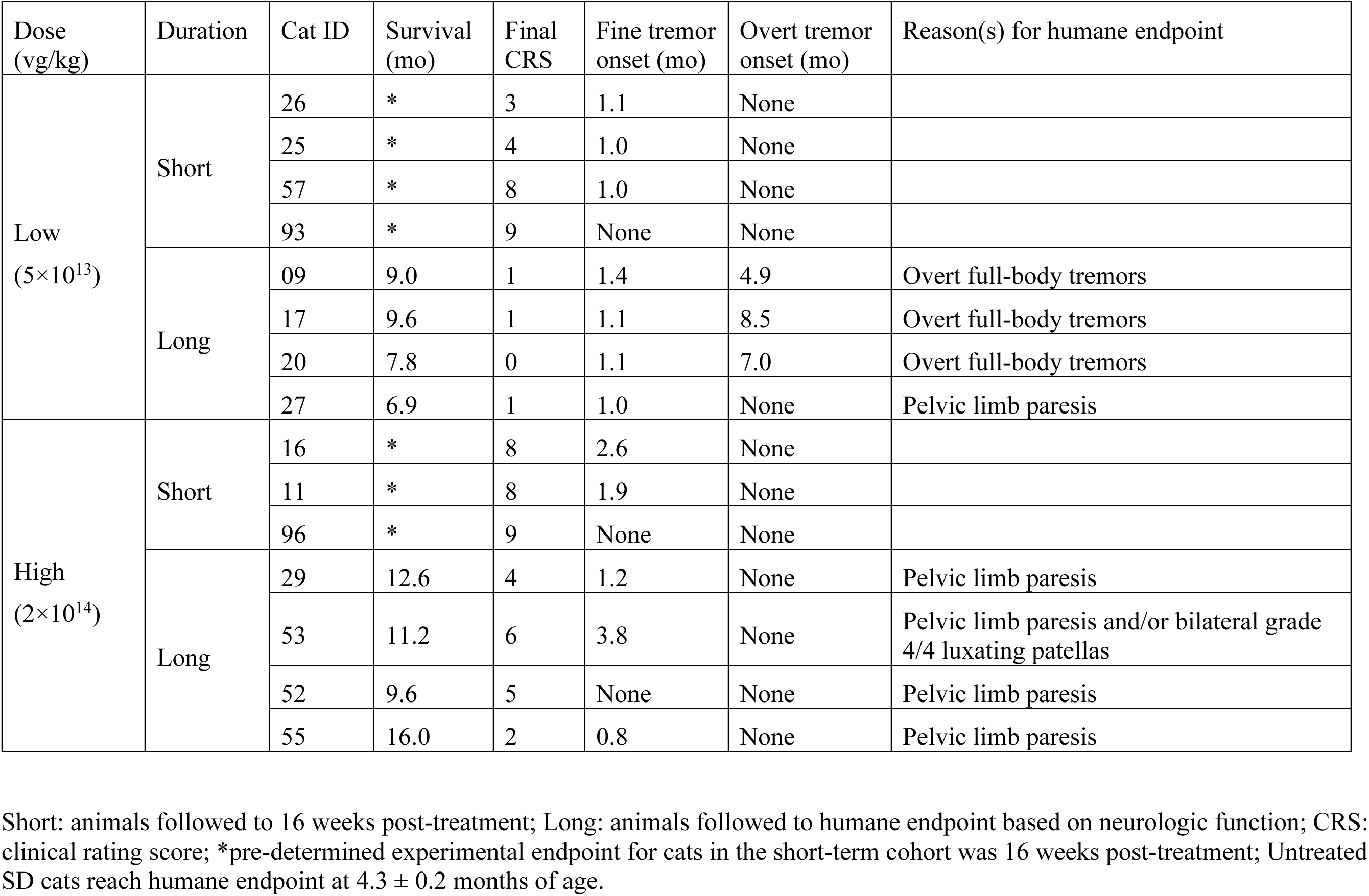
AAV-treated SD cats and their clinical outcomes.

| Dose (vg/kg) | Duration | Cat ID | Survival (mo) | Final CRS | Fine tremor onset (mo) | Overt tremor onset (mo) | Reason(s) for humane endpoint |
| --- | --- | --- | --- | --- | --- | --- | --- |
| Low ($5 \times 10^{13}$) | Short | 26 | * | 3 | 1.1 | None | |
|  |  | 25 | * | 4 | 1.0 | None |  |
|  |  | 57 | * | 8 | 1.0 | None |  |
|  |  | 93 | * | 9 | None | None |  |
|  | Long | 09 | 9.0 | 1 | 1.4 | 4.9 | Overt full-body tremors |
|  |  | 17 | 9.6 | 1 | 1.1 | 8.5 | Overt full-body tremors |
|  |  | 20 | 7.8 | 0 | 1.1 | 7.0 | Overt full-body tremors |
|  |  | 27 | 6.9 | 1 | 1.0 | None | Pelvic limb paresis |
| High ($2 \times 10^{14}$) | Short | 16 | * | 8 | 2.6 | None | |
|  |  | 11 | * | 8 | 1.9 | None |  |
|  |  | 96 | * | 9 | None | None |  |
|  | Long | 29 | 12.6 | 4 | 1.2 | None | Pelvic limb paresis |
|  |  | 53 | 11.2 | 6 | 3.8 | None | Pelvic limb paresis and/or bilateral grade 4/4 luxating patellas |
|  |  | 52 | 9.6 | 5 | None | None | Pelvic limb paresis |
|  |  | 55 | 16.0 | 2 | 0.8 | None | Pelvic limb paresis |

Short: animals followed to 16 weeks post-treatment; Long: animals followed to humane endpoint based on neurologic function; CRS: clinical rating score; \*pre-determined experimental endpoint for cats in the short-term cohort was 16 weeks post-treatment; Untreated SD cats reach humane endpoint at $4.3 \pm 0.2$ months of age.

### Ultrasound elastography

Ultrasound elastography measures the elastic properties of soft tissue through the propagation of shear waves. A loss of elasticity, or increase in tissue stiffness, is reported in common conditions such as breast cancer and liver fibrosis. We hypothesized that an increase in storage material in SD tissues would decrease elasticity (or increase stiffness).

Two common readouts from elastography are wave speed and elasticity, which in the liver of SD cats were significantly reduced compared to normal. AAV treatment led to a transient, dose-dependent pattern of improvement. As shown in Fig. 2, wave speed and elasticity approached normal in the liver of SD cats treated with the high dose until 7 months of age but returned to untreated levels at later time points. SD cats treated with the low dose had wave speed and elasticity reductions similar to untreated SD cats at all time points. The spleen and kidney of untreated SD cats had wave speed and elasticity reductions that did not reach statistical significance compared to normal cats. Mean values in pancreas and skeletal muscle were similar to normal. Neither AAV dose appeared to have any effect on elastography metrics in spleen, kidney, pancreas or skeletal muscle (Fig S2).

**Figure 2.**
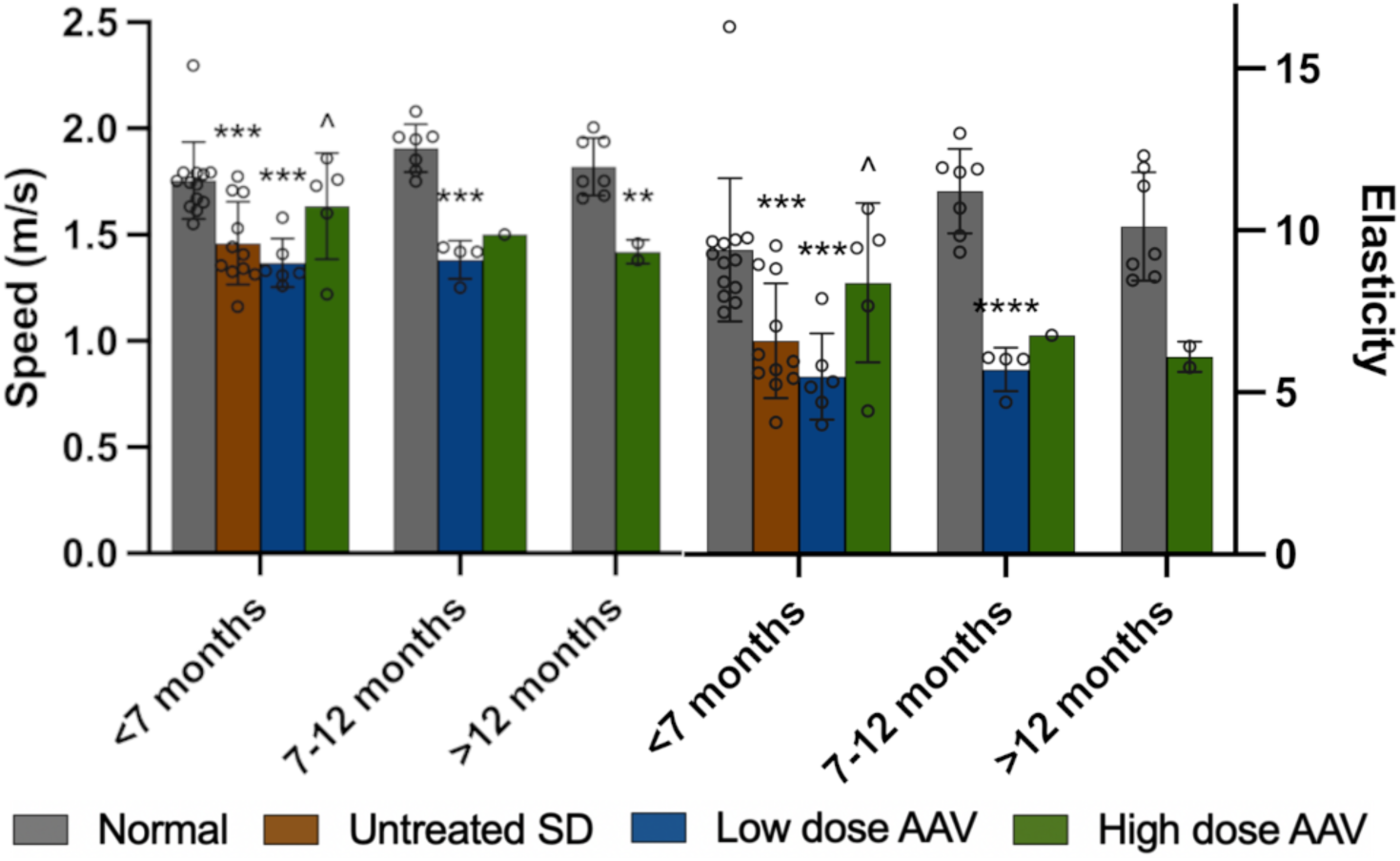
Liver stiffness in SD cats is improved with high dose AAV treatment. Shear wave ultrasound elastography demonstrated significant decreases in speed and elasticity at all ages of untreated and low dose treated SD cats compared to normal liver. In SD cats treated with the high dose, wave speed and elasticity returned to near-normal levels until 7 months of age, after which they decreased to untreated levels. *0.05>p>0.01, **0.01>p>0.001, ***0.001<p<0.0001, ****p<0.00001 compared to age-matched normal cats; ^p<0.05 versus low dose at same age.

### 7T MR images and MRS metabolites

T2-weighted MR images acquired with a 7T (Tesla) scanner demonstrate that AAV treatment partially restores gray/white matter distinction and minimizes brain atrophy (Figure 3A). In T2 images from normal cats, white matter is darker than (hypointense to) gray matter in the cerebrum and cerebellum. For example, the corona radiata is clearly differentiated from surrounding gray matter of the parietal cortex and the white matter of the deep cerebellar nuclei are distinct from overlying cerebellar cortex. In untreated SD cats, an increase in white matter signal intensity and decrease in gray matter signal intensity causes the normally clear gray-white matter boundaries to become isointense. In addition, the hyperintense CSF signal surrounding the brain and gyri becomes wider in SD cats, as the brain parenchyma atrophies and CSF fills the resulting space. Images of AAV-treated cats show partial restoration of the gray/white matter boundaries in the cerebrum / cerebellum, especially in the high-dose, short-term cohort imaged 16 weeks after treatment. Similarly, the most pronounced preservation of CSF volumes was apparent in the high-dose, short-term cohort. As AAV-treated cats in the low-dose and high-dose cohorts neared humane endpoint, deterioration of brain MRI architecture (i.e., isointensity of gray-white matter boundaries and increased CSF volumes) became more similar to untreated SD cats (Figure 3A). Thus, MRI architecture of the brain correlated well with clinical disease in AAV-treated and untreated SD cats over time.

**Figure 3.**
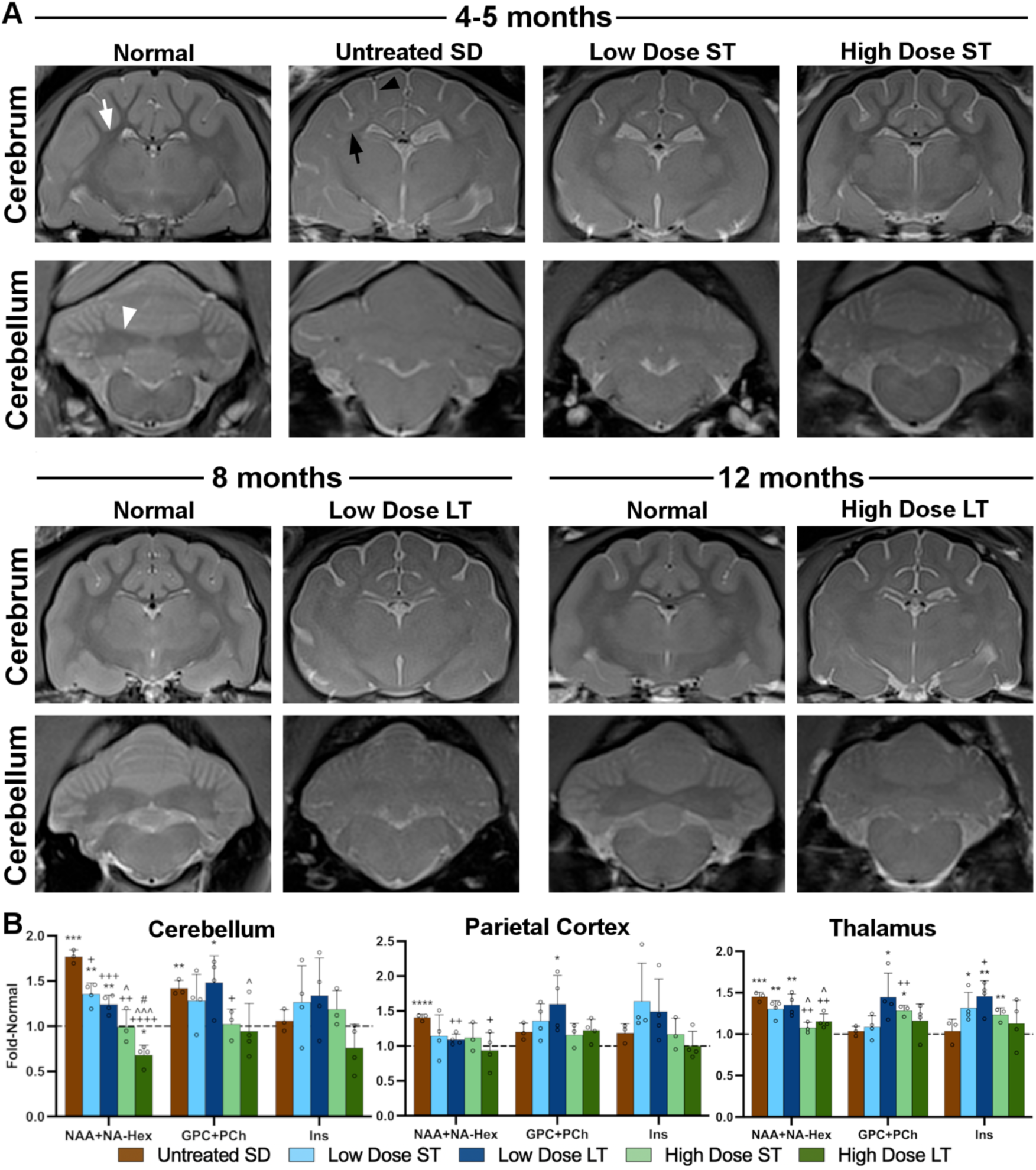
AAV treatment partially normalizes 7T MR images and MRS metabolites. (A) T2-weighted anatomical images. In normal cats, white matter is darker than, or hypointense to, gray matter in the corona radiata of the cerebrum (white arrow) and the DCN of the cerebellum (white arrowhead). The CSF signal (bright) around the meninges is minimal in normal cats. In untreated SD cats at humane endpoint, the DCN are isointense to (same shade as) surrounding gray matter and the corona radiata (black arrow) are hyperintense to (lighter than) gray matter due to hypomyelination and storage in neuron cell bodies. The increased amount of CSF signal (black arrowhead) in untreated SD cats is especially apparent surrounding the gyri and is attributed to generalized atrophy. Pathological MRI changes are mitigated by AAV treatment, with the most prominent preservation of brain architecture in the high-dose, short-term cohort. (B) MRS of 3 brain voxels: cerebellum, parietal cortex, thalamus. The spectroscopic peak of the neurometabolic marker NAA, which merges with the NA-Hex storage product in SD, is significantly increased in all regions of SD cat brains and normalized by the high dose of AAV in the cerebellum and thalamus. GPC+PCh, a measure of membrane turnover that correlates with demyelination in this model, is significantly increased in the cerebellum of untreated SD cats and normalized in the high-dose cohort at both short- and long-term timepoints. Myoinositol (Ins) is unchanged in untreated and treated SD cats, though variability is high in some groups. ST: Short-term, LT: Long-term. *0.05>p>0.01, **0.01>p>0.001, ***0.001<p<0.0001, ****p<0.00001 vs. age-matched normal cats; ^+^0.05>p>0.01, ^++^0.01>p>0.001, ^+++^0.001<p<0.0001, ^++++^p<0.00001 vs. untreated SD cats; ^0.05>p>0.01, ^^^0.001<p<0.0001 vs. low dose at same age. ^#^p<0.05 vs. short-term at same dose. Dashed horizontal line represents levels in age-matched normal cat brains, to which all SD data were standardized.

MRS data suggest that the cerebellum is more impacted in untreated SD cats than the parietal cortex or thalamus, with significant increases in both NAA+NA-Hex (N-acetylaspartate + N-acetylhexosamine) and GPC+PCh (glycerophosphocholine + phosphocholine) (Figure 3B). While NAA reflects neuronal health and is decreased in many neurodegenerative storage diseases, NA-Hex is a storage product that cannot be resolved spectroscopically from NAA. Thus, the overlapping spectra of NAA+NA-Hex serve as a biomarker of storage in the feline SD brain, with elevations also occurring in the parietal cortex and thalamus. In contrast, the parietal cortex and thalamus of untreated SD cats have normal levels of GPC+PCh, an indicator of membrane turnover that often correlates with demyelination. Myoinositol (Ins), thought to be an indicator of neuroinflammation, remains unchanged in all 3 (Figure 3B).

The high dose of AAV normalized NAA+NA-Hex in all 3 brain voxels, in both short-term and long-term cohorts. A dose effect was noted in the cerebellum and thalamus of SD cats, with the low dose yielding NAA+NA-Hex levels that were intermediate to the untreated and high dose cohorts. The high dose of AAV reduced GPC+PCh levels to normal in the cerebellum, though its effect in the parietal cortex and thalamus was less conclusive due to insignificant elevations of GPC+PCh in these voxels from untreated SD cats (Figure 3B).

### CSF biomarkers of pathology

CSF levels of aspartate aminotransferase (AST) and lactate dehydrogenase (LDH), indicators of cell damage within the CNS, were significantly increased in untreated SD cats at humane endpoint (4-5 months of age) (Figure 4A, B). AAV-treated SD cats had less AST and LDH at 5 months of age than untreated SD cats, with the high dose being more effective at reducing these levels (Figure 4A, B). Specific activity of all isozymes of Hex (Total Hex) was decreased to background levels (6.85% of normal) in untreated SD cats and restored to near-normal or above-normal levels in cats treated with both doses across ages (Figure 4C).

**Figure 4.**
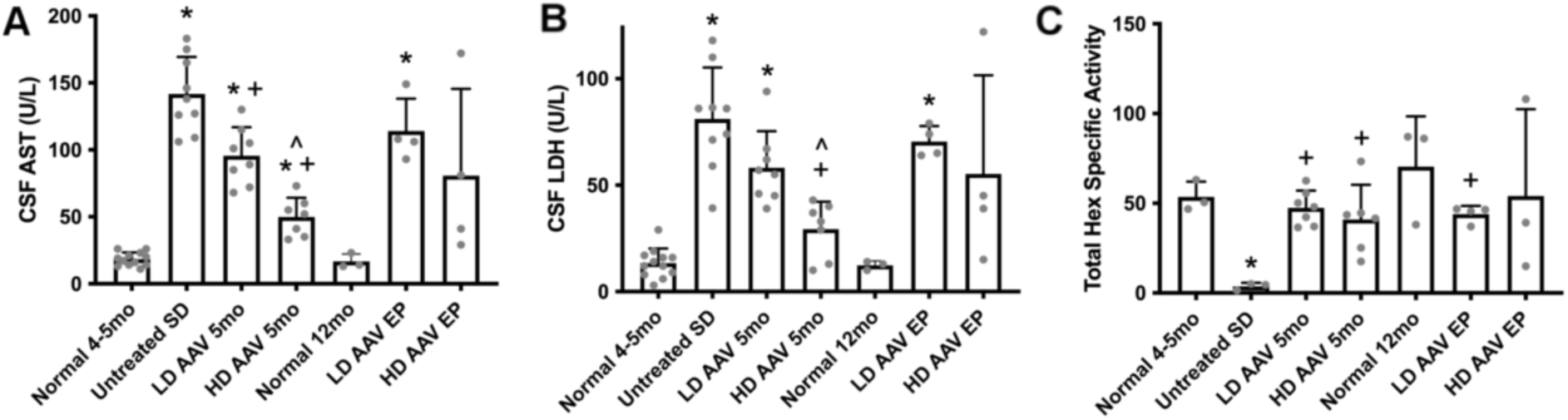
AAV treatment reduces CSF markers of cell damage and increases Hex activity. (A, B) CSF levels of aspartate aminotransferase (AST) and lactate dehydrogenase (LDH) are significantly increased in SD cats, indicating cell damage within the central nervous system. AST and LDH are significantly reduced in cats treated with the high AAV dose at the short-term time point (5 months of age). AST and LDH rose between the short- and long-term time points as clinical disease progressed. (C) Specific activity of all hexosaminidase isozymes (Total Hex) is significantly decreased to background levels in the CSF of untreated SD cats. AAV treatment normalizes Total Hex levels across doses and age groups. ST, short-term; LT, long-term. *p<0.05 versus age-matched normal cats, ^+^p<0.05 versus untreated SD cats, ^p<0.05 versus low dose at same age.

### HexA activity in CNS

Specific activity levels of HexA, the isozyme responsible for degrading GM2 ganglioside in humans, were improved in AAV-treated cohorts, with variability dependent on CNS region and dose. In general, HexA activity remained near baseline (untreated) levels in the rostral portion of the forebrain, with a slight, dose-dependent increase in the caudal forebrain (Figure 5A, blocks A-F). In the hindbrain and spinal cord of treated SD cats, HexA activity increased significantly to as high as 0.7- and 1.2-fold normal, respectively (Figure 5A, blocks G-I and J-P). A dosage effect was noted in the hindbrain and spinal cord, where mean HexA activity in the high dose cohort was 6.13- and 6.73- fold that of the low dose cohort, respectively. Based on restoration of HexA activity, the spinal cord of the high dose cohort was treated most effectively of all CNS regions, with mean activity ranging from 0.2 to 1.0 fold normal HexA in the long-term cohort (Figure 5). Similar results were obtained when using a substrate that measures the activity of all Hex isozymes (MUG; Figure S3).

**Figure 5.**
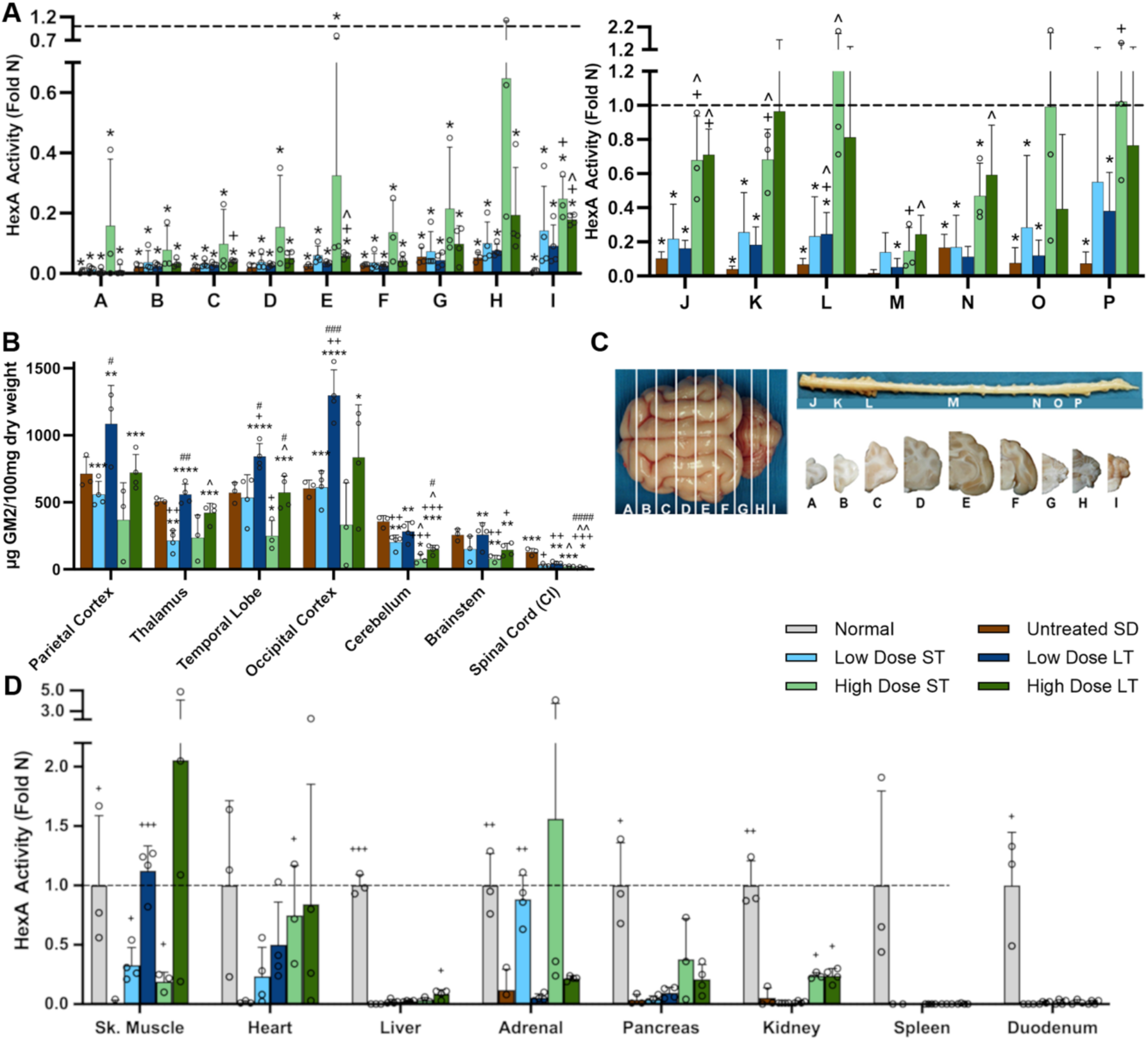
Hex activity increases and GM2 ganglioside storage decreases with AAV treatment. (A) HexA activity (fold normal, or Fold N) in the CNS increases with AAV dose, with the cerebellum, cervical spinal cord, and lumbar spinal cord showing the best response. (B) GM2 ganglioside in untreated SD cats was significantly increased above normal (trace) levels in all CNS regions except spinal cord. AAV treatment reduced GM2 levels in certain regions of the CNS (depending on dose and time point) but only restored it to normal levels in the cervical intumescence (CI) of the spinal cord. (C) Brain and spinal cord block designations. Dorsal surface of the brain is shown on the left, with corresponding transverse hemi-sections on the right (below spinal cord). (D) HexA activity in peripheral tissues, including the legend for panels A-D. *p<0.05, **0.01>p>0.001, ***0.001<p<0.0001, ****p<0.00001 vs. age-matched normal cats; ^+^p<0.05, ^++^0.01>p>0.001, ^+++^0.001<p<0.0001, ^++++^p<0.00001 vs. untreated SD cats; ^0.05>p>0.01, ^^0.01>p>0.001, ^^^0.001<p<0.0001 vs. low dose at same age. ^#^p<0.05, ^##^0.01>p>0.001, ^###^0.001<p<0.0001, ^####^p<0.00001 vs. short term at same dose.

HexA activity levels in untreated SD cats were ≤ 10% of normal across all CNS sections, with the exception of the thoracolumbar region (block N) at 17%. The brain activity of lysosomal enzymes other than Hex was above normal, a well-known phenomenon thought to represent a compensatory mechanism when one lysosomal enzyme is missing. For example, lysosomal enzymes ß-galactosidase (Bgal) and α-mannosidase (Mann) had high levels of activity in untreated SD cats that were reduced toward normal in effectively treated regions of the brain and spinal cord (Fig S3). Secondary lysosomal enzyme activity has been proposed as a biomarker of disease progression in SD and similar storage disorders.

### GM2 ganglioside storage

GM2 ganglioside was elevated throughout the CNS of untreated SD cats compared to normal cats, which have only trace amounts of GM2 (Figure 5B). In SD cats treated with the high dose of AAV, mean levels of GM2 were slightly or significantly reduced throughout the CNS at 16 weeks post-treatment. At humane endpoint, GM2 levels returned to those of untreated SD cats in the cerebrum but remained reduced in the cerebellum, brainstem and spinal cord, mirroring the higher HexA activity in those regions. GM2 levels in the CNS of the low-dose cohort were more variable, with significant reduction in the thalamus at 16 weeks post-treatment but no reduction in other regions. At humane endpoint of the low-dose cohort, GM2 again remained at untreated levels in most regions and even was significantly elevated in the occipital cortex to more than twice that of untreated SD cats (Figure 5B). Levels of sialic acid, a constituent of GM2 ganglioside, generally mimicked those of GM2 in the CNS of untreated and treated SD cats (Fig S4).

### HexA activity in peripheral tissues

HexA activity in peripheral tissues of AAV-treated SD cats varied according to tissue type and duration of follow-up. For example, normal to above-normal HexA activity was measured in the heart, skeletal muscle and adrenal gland while only background activity was produced in the spleen and duodenum (Fig. 5D). Dose and treatment duration influenced activity in some tissues such as heart, where HexA ranged from a low of 0.19 fold normal (low dose, short term) to a high of 0.84 fold normal (high dose, long term). Activity in the skeletal muscle was only 0.2 – 0.3 fold normal in short term cohorts but increased to 1.1 – 2.1 fold normal in animals followed long term, suggesting that HexA activity increased slowly over many months. In general, the highest levels of HexA were produced by tissues with the highest number of vector genomes. However, a surprising exception was noted in liver, which had low HexA activity that never exceeded 0.09 fold normal at any dose or time point, even though liver had the highest number of vector genomes of all tissues (Fig. S7). HexA activity in the adrenal gland varied across cohorts but approached normal levels in the low dose, short term cohort. Other tissues, such as kidney and pancreas, had intermediate levels of HexA activity ranging from ∼0.2 – 0.4 fold normal in high dose-treated groups (Fig. 5D).

### Neuroinflammation

Gliosis is a known feature of untreated SD cats as shown by abnormally high levels of the astrocytic marker, GFAP (glial fibrillary acidic protein) and the microglial marker, Iba1 (ionized calcium-binding adaptor molecule 1). For example, GFAP pixel density was ∼12 fold above normal in gray matter of the parietal cortex in untreated SD cats (Fig. 6A). Though Iba1 staining increases were less pronounced, microglial activation was clear through changes from a ramified morphology in the normal cat brain to an ameboid morphology in the SD cat brain. That is, normal microglia were ramified with a small cell body and long, branching processes compared to the ameboid morphology of SD microglia, with an enlarged cell body and shortened, blunt processes (Figure 6B).

**Figure 6.**
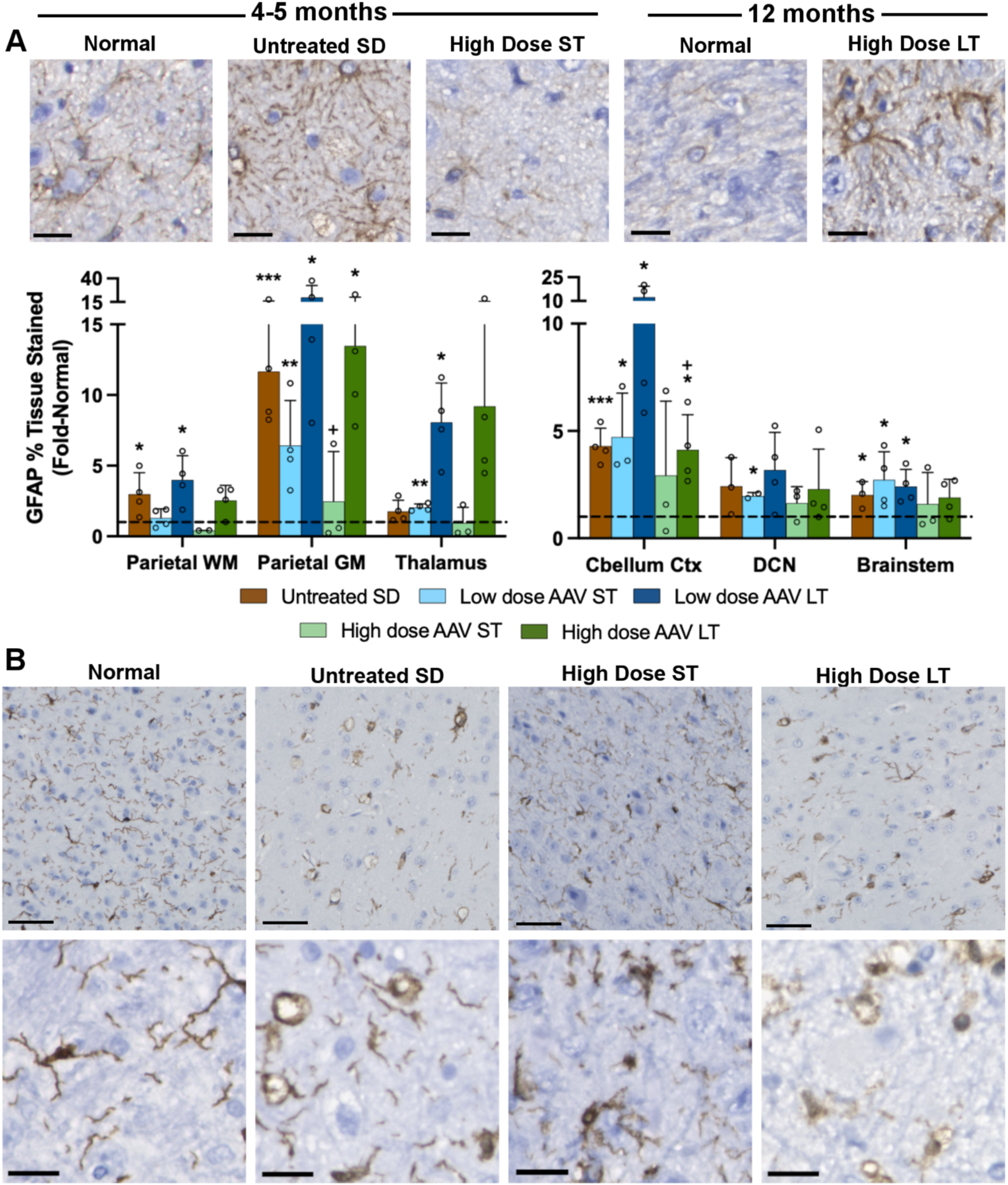
Astrocyte and microglia pathology is transiently normalized after AAV treatment. SD cats treated with the high dose were evaluated short-term (16 weeks post-treatment or ∼5 months of age) and long-term (humane endpoint at ∼12 months of age). Brain sections were stained for (A) GFAP (astrocyte marker) or (B) Iba1 (microglia marker). Micrographs at the top of each panel show representative staining in the thalamus and demonstrate normalization of glial cell morphology in the short-term cohort but not in the long-term cohort. Graphs depict the percentage of positive staining for each marker standardized to that of normal cats (represented by dashed horizontal line). Scale bars: 20 µm. ST: short-term, LT: long-term, WM: white matter, GM: gray matter, DCN: deep cerebellar nuclei. *p<0.05, **0.01>p>0.001, ***0.001<p<0.0001 vs. age-matched normal cats; ^+^p<0.05 vs. untreated SD cats .

With AAV treatment, glial cell morphology was largely normalized in animals of the high-dose, short-term cohort (16 weeks post-injection, or ∼5 months of age). For example, astrocyte morphology in treated SD cats was similar to that of normal cats, but GFAP stain intensity and thickness of the cell body / processes were increased in untreated SD cats (Fig. 6A). Also, ameboid microglia were prominent in untreated SD cats but ramified microglia predominated in normal and AAV-treated SD cats in the short-term cohort (Fig. 6B). In contrast, animals of the high-dose, long-term cohort (followed to the humane endpoint of 12.4±2.7 months of age) had astrocytic and microglial morphology that resembled untreated SD cats, suggesting that glial cell pathology was reversed only temporarily by AAV treatment. Such findings are consistent with a previous study in which SD cats treated by intracranial delivery of AAV had transient normalization of glial cell pathology (*52*).

### Oligodendrocytes and myelin

The number of oligodendrocytes labeled by Olig2 was reduced to around 50% normal in the parietal cortex, thalamus, and cerebellar cortex, and around 75% in the DCN and brainstem (Figure 7). In the parietal cortex and thalamus, AAV treatment partially normalized oligodendrocyte levels in a dose-dependent fashion. In the cerebellar cortex, DCN, and brainstem, oligodendrocyte numbers were closer to normal with both dose groups, with long-term cats having above-normal levels of oligodendrocytes in the brainstem. The high variance of the LFB staining makes it difficult to draw concrete conclusions, but in general the most affected areas in untreated SD cats are the parietal cortex, thalamus, and cerebellar cortex, with dose-dependent partial normalization. LFB staining in the DCN and brainstem was at approximately normal levels across untreated and all AAV-treated cats.

**Figure 7.**
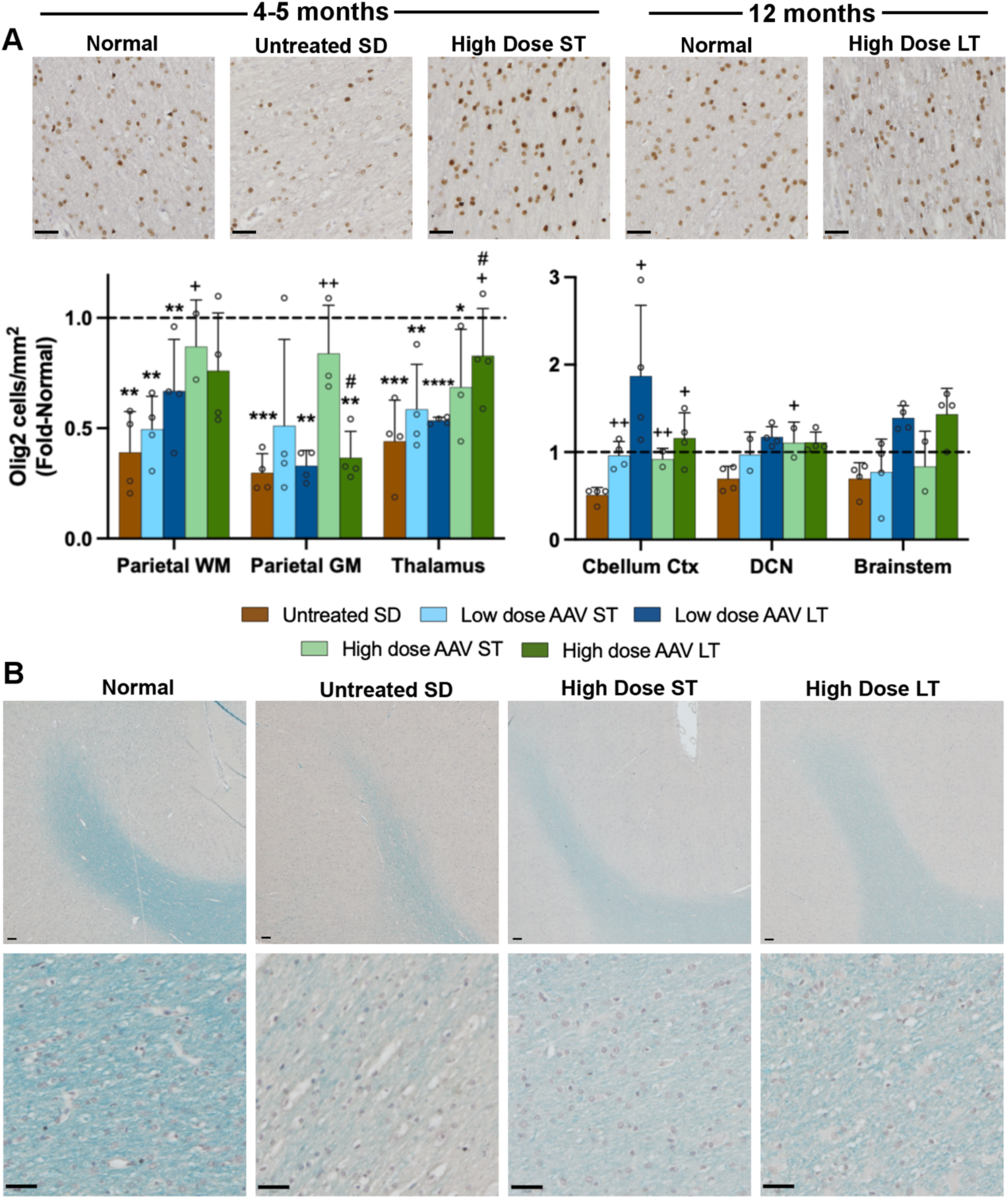
Myelin deficits in SD cats are partially normalized through AAV treatment. (A) Oligodendrocytes staining positive for Olig2 and (B) percentage of tissue staining positive for Luxol Fast Blue (LFB) are below normal levels in SD cats and partially restored in AAV-treated cats. Micrographs at 20x are from the parietal cortex white matter of representative cats. Scale bar: 20 µm. ST: short-term, LT: long-term, WM: white matter, GM: gray matter, DCN: deep cerebellar nuclei. *p<0.05, **0.01>p>0.001, ***0.001<p<0.0001, ****p<0.00001 vs. age-matched normal cats; ^+^p<0.05, ^++^0.01>p>0.001 vs. untreated SD cats; ^#^p<0.05 vs. short term at same dose.

### Histopathology

Histopathological changes for untreated SD cats at humane endpoint include moderate-marked swelling of neurons due to storage material, foamy macrophages surrounding vessels, mild-moderate gliosis, and dilated myelin sheaths with Gitter cells in white matter tracts throughout the CNS (Figure S5). Cats in the high dose short-term AAV cohort had the fewest lesions overall, with only mild, rare neuronal storage and white matter deficits only evident in the cerebellum. The low dose short-term AAV cohort demonstrated some improvement in severity of CNS lesions, while the long-term cohorts of both doses had severe lesions on the level of untreated SD cats. In the liver, storage lesions (vacuolation) in hepatocytes were marked in untreated SD cats (Figure S5). Vacuolation remained apparent in most AAV-treated cohorts, consistent with unexpectedly low HexA activity in the liver of treated cats. However, vacuoles appeared to be smaller in the high-dose cohort versus the low-dose cohort. The greatest improvement in hepatocyte pathology occurred in the high dose long-term cohort, which had the highest level of HexA activity (0.09 fold, or 9% of, normal).

### AAV vector biodistribution and serum antibody titers

AAV vector was present throughout the CNS and peripheral tissues in treated animals, with the high dose and short-term groups generally having the highest levels (Figure S6). In the CNS, vector levels were highest in the cerebral cortex (0.050-9.658 vg/kg), with the cerebellum, brainstem and spinal cord having similarly low levels (0.004-1.02 vg/kg). The levels of vector were more dependent on dose than time post-treatment, with the high dose cohorts having higher vector levels than low dose cohorts within the CNS. Similar dose-dependent differences in vector copies were also noted in the PNS tissues (sciatic nerve, cervical and lumbar DRGs), while other peripheral organs were relatively homogeneous among cohorts. Among the peripheral tissue, the liver had the highest level of vector. Tissues with vector <0.2 vg/kg were kidney, spleen, duodenum, and pancreas. Post-treatment, all cats developed serum antibody titers to AAV9 ranging from 1:1000 - 1:256,000(Fig S7). All cats for whom pre-treatment serum was collected (n = 13) had undetectable serum titers below 1:8 dilution.

## DISCUSSION

Enzyme replacement as a treatment for storage diseases has been contemplated since the 1960-70s, when stored substrate was eliminated from mucopolysaccharidosis fibroblasts by “corrective factors” from cell culture supernatant, cell homogenates or normal body fluids (*38*). Similar results were reported for other storage diseases. In most cases, the corrective factors proved to be lysosomal hydrolases that could be endocytosed by enzyme-deficient cells and restore normal metabolism. Since that time, enormous efforts have been made to deliver functional enzyme to patients. For conditions such as Gaucher disease type I, with primary pathology outside of the CNS, enzyme replacement therapy has been quite successful. Enzyme replacement is a far greater challenge for the ∼70% of storage disorders that are neuropathic, like Tay-Sachs and Sandhoff diseases, in which the BBB blocks functional enzyme from entering the parenchyma of the brain and spinal cord. Thus, effective CNS delivery mechanisms may be the next great advance in treating storage disorders, akin to the discovery of “corrective factors” in the 1960s.

Such is the hope for AAV gene therapy, in which the vector is both a means of enzyme replacement and a mechanism for delivery across the BBB. In the current study, the IV-delivered bicistronic AAV9 vector clearly improved lifespan and quality of life in SD cats in a dose-dependent manner. Longevity almost doubled with the low dose and tripled with the high dose, and AAV-treated cats maintained a higher weight and CRS than their untreated counterparts. Similar to previous studies of AAV treatment in SD cats (*9*), by far the most dramatic clinical improvement was the reduction of severe overt tremors, presumed to be primarily cerebellar in origin. Overt tremors only developed in 3 of 15 AAV-treated cats in the current study (all in the low dose cohort), with the age of onset delayed by 5 months compared to untreated SD cats.

MR images reveal partial correction of brain atrophy and gray/white matter distinction with IV treatment, similar to results from intracranial AAV therapy in SD (*9*) and GM1 (*39*) cats. Isointensity of gray/white matter boundaries is a feature typical of human GM2 (*13*) and GM1 gangliosidoses (*40*), and MR-based monitoring of brain changes will provide important metrics of disease progression in clinical trials. Further evidence of efficacy in IV-treated SD cats was revealed by brain metabolite levels measured with MRS. GPC+PCh, an indicator of membrane turnover that correlates with myelin abnormalities, was significantly elevated in the cerebellum of SD cats and returned to normal after IV treatment with the full dose. As a marker of neuronal health, NAA levels are abnormally low in GM1 cats (*37*) and other CNS diseases such as multiple sclerosis (*41*) and intra-cerebral hemorrhage (*42*). However, in SD, the NAA peak cannot be fully resolved from a storage product, NA-Hex, so the combined NAA+NA-Hex peak actually serves as an *in vivo* marker of brain pathology. Elevations of NAA+NA-Hex in the brain of SD cats were reduced to normal after high-dose IV treatment at both short- and long-term follow-up.

CSF analysis provides another medium for *in vivo* assessment of SD pathology and AAV treatment efficacy. AST and LDH are ubiquitously expressed enzymes that are released from cells following plasma membrane damage. High levels in the CNS are well-documented in a variety of neurological diseases including human TSD (*43, 44*), canine GM1 gangliosidosis (*45*), and human meningitis (*46*). Similar to intracranial AAV treatment in SD (*47*) and GM1 (*48*) cats, IV treatment of SD cats in this study resulted in normalization of AST and LDH. Also measurable in the CSF was Hex specific activity, which is minimal in untreated SD cats. Hex activity increased to normal levels in both doses of AAV-treated cats.

Though survival in IV-treated cats is promising, reaching 3-fold longer than untreated SD cats, it did not equal the quadrupling of lifespan achieved by intracranial treatment. In fact, intracranial injection corrected CNS pathology so effectively that approximately half of the treated SD cats died of peripheral organ disease (*9*). Gross enlargement of the stomach, urinary bladder and colon found frequently in long-term follow-up after intracranial treatment, were not an important factor in the demise of IV-treated cats. Thus, it is likely that IV administration corrected some of the peripheral organ pathology that was not treated effectively by intracranial injection. Further confirmation of this hypothesis comes from ultrasound shear wave elastography, which demonstrated significant liver stiffness in SD cats that was corrected by the high dose of AAV treatment. This noninvasive method is increasingly accepted for monitoring of conditions such as liver fibrosis (*31*), intervertebral disc pathology (*32*), and cardiac disease (*33*) in human patients, and has recently been adapted for use in feline clinical medicine (*34–36*). Elastography shows promise as an *in vivo* method to monitor peripheral organ disease in ongoing or future clinical trials.

A peripheral disease manifestation that was not effectively addressed by either intracranial or IV routes was spinal cord compression, evident upon necropsy of most AAV-treated cats. Clinically, these cats often displayed pelvic limb paresis or paralysis that led to the predetermined humane endpoint (inability to stand). Bony abnormalities have been recorded in radiographs and 3T MR images in the cervical spine and coxofemoral regions in SD cats, though the pathogenesis of these changes is unknown (*29*). These clinical signs are also observed in human patients with mucopolysaccharidosis (MPS), a family of LSDs that primarily affects connective tissue (*30*). In animal models of MPS, spinal cord compression has been refractive to treatment, responding only to enzyme replacement at high doses and very early ages (*49*). Future gene therapy studies should focus on strategies to effectively reduce spinal cord compression, which will benefit patients with a variety of lysosomal storage disorders.

Improvement in post-mortem Hex activity was greatest in the spinal cord, with most cats achieving therapeutic levels of 20% of normal, the threshold at which humans are clinically asymptomatic. IV delivery did not result in the same levels of Hex that intracranial treatment achieved in SD cats, in which all brain regions had above-normal levels of activity (*27, 28*), but faithfully recapitulates the Hex activity seen in SD mice with IV delivery of this bicistronic vector (*26*). Similar to mouse studies, the spinal cord appears to be the best-transduced area, with mixed results in the brainstem, cerebellum, and cerebral cortex (*26, 50*). The specific activity of HexA and Total Hex is increased in AAV-treated cats and inversely correlated with storage of GM2 ganglioside, total sialic acid, and the activity of compensatory hydrolases (Bgal and Mann), consistent with previous AAV studies in cats and mice(*26, 28, 51, 52*). Notably, cats in the high dose long-term group often had lower levels of storage than cats in the low dose group, despite the fact that they lived 4 months longer on average.

Activation of astrocytes and microglia, hallmarks of neuroinflammation, was mitigated 16 weeks after AAV treatment but not in longer-lived cats. GFAP-positive staining was increased in untreated SD cats and long-term AAV-treated cats, coupled with the activated astrocyte morphology of thickened cellular processes (*53*). Microglia followed a similar pattern, with increases in quantity and activated morphology (ameboid, with large cell bodies and short, thick processes), in untreated SD and long-term AAV treated cats. These results are consistent with previously published studies in SD (*54*) and GM1 (*37, 41*) cats, in which the increases of Iba-1 and GFAP staining, respectively, in untreated cats were reduced by intracranial AAV injection. A likely explanation is that AAV treatment is effective at reducing neuroinflammation initially, but inciting causes of neuroinflammation remain unresolved, leading to accumulation in longer-lived cats. Cats in the high dose group still had lower levels of inflammation than the low dose group, despite the fact that they lived 4 months longer on average, supporting the hypothesis of dose-dependent efficacy.

Oligodendrocyte number and myelin are below normal levels in most regions measured, with partial normalization through AAV treatment. In untreated GM1 cats, a reduction in Olig2-positive cell numbers was not observed in the 6 regions examined (striatum, thalamus, parietal cortex, temporal lobe, occipital cortex, and cerebellum) (*37*), in contrast to the current study in which it was observed in all regions. A biological explanation for this difference in myelin pathology between two otherwise very similar pathological processes is not obvious, though perhaps differences in methodology or severity of clinical disease could be the cause. This could be attributed to pathology within existing oligodendrocytes being compensated for post-treatment by differentiation of oligodendrocyte precursor cells into new mature oligodendrocytes. Myelin loss, as determined by a decrease in LFB staining, matched GM1 cats in that there was extreme loss in the parietal cortex and cerebellum and less severe loss in the thalamus (*37*).

Mild pathology in the brainstem of untreated SD cats is not limited to mild decreases in myelin and oligodendrocyte number, as neuroinflammation and neuronal loss are least severe in the brainstem as well. In addition, AAV treatment may have a more dramatic effect in the brainstem than other regions, since neuroinflammation does not appear to increase in long-term treated cats, and oligodendrocytes increase to above-normal levels. When considered with the Hex activity and storage data, the brainstem appears to be the best-treated region in the brain, though this could be due partly to milder disease in the brainstem at the outset.

The variability in longevity, clinical signs, and Hex activity in the long-term groups (of both doses) supports the previously established hypothesis that AAV treatment effectively shifts the phenotype in SD cats from one that imitates the infantile form to one that imitates a later-onset less severe form. Patients with the infantile form of GM2 gangliosidosis possess mutations that render Hex either unable to form or dysfunctional, while juvenile or adult-onset patients possess mutations that allow for some Hex to form properly and function, albeit at levels much below normal. By introducing functional copies of *HEXA* and *HEXB* cDNAs to SD cats and causing modest increases in Hex activity, we created a crude representation of a delayed-onset phenotype. The longer the treated cats live, the more heterogenous their lifespan and other clinical signs, which recapitulates human juvenile and adult-onset GM2 gangliosidosis. Therefore, AAV-treated cats provide valuable information not only about response to treatment, but also about late-onset subtypes of SD and TSD.

This parallel between the AAV-treated feline model and later-onset GM2 patients offers insight into the improvement of treatment success. Though all *in vivo* metrics (elastography, MRI and MRS, CSF biomarkers) demonstrated some degree of normalization across all age groups, post-mortem metrics (storage, inflammation, myelin deficits and histopathological lesions) followed a pattern of improvement in the short-term cohort that waned in the long-term cohort. Such transient effect may suggest that a pathological cascade is set in motion before the age of treatment that is slowed but not eliminated by AAV gene therapy. Though a subsequent administration of AAV vector is one solution, the development of anti-capsid titers in treated cats suggests that re-administration of vector will be ineffective and even dangerous. Subsequent improvements in AAV treatment of GM2 gangliosidosis are therefore likely to come from early intervention and enhancement of vector design.

## MATERIALS AND METHODS

### Study Design

Untreated SD cats were randomly assigned to low or high dose treatment groups, with at least one male and female in each group. The first cats treated for each dose were assigned to the long-term treatment group to allow the study to conclude in a timely manner. A sample size of 4 cats per group was predetermined, but limitations in vector manufacturing resulted in one treatment group with n=3 (high dose, short-term).

### AAV Vector Preparation

Preparation of the AAV9 bicistronic vector was performed similarly to previously published methods of AAV9-Bic in mice (*26*), except that feline HEXA and HEXB cDNAs were used in this study. The vector includes a centrally located, bidirectional enhancer/promoter complex consisting of a cytomegalovirus (CMV) enhancer flanked by 2 chicken β-actin promoters in opposite directions. The low and high doses were 5E13 vector genomes/kg and 2E14 vector genomes/kg, respectively.

### Animals and AAV Treatment

A breeding colony of SD cats is maintained at the Scott-Ritchey Research Center at Auburn University, and all experiments in this study were approved by its Institutional Animal Care and Use Committee. Affected SD cats were treated between 0.77 and 1.47 months of age (mean 1.10±0.15) and 0.38 and 0.70 kg (mean 0.49±0.15). An intravenous catheter was placed in the cephalic vein and flushed with saline. Vector was delivered slowly over a period of 1-2 minutes via the catheter, which was subsequently flushed with saline. Cats were monitored for at least 3 hours post-treatment for any adverse reactions. Euthanasia was performed with cardiac injection of pentobarbital and animals were subsequently perfused with heparinized saline. AAV-treated cats were euthanized 16 weeks post treatment (low dose n=4, high dose n=3) or followed until they reached the humane endpoint of inability to stand (low dose n=4, high dose n=4). Controls were age-matched untreated normal cats (n≥3) and untreated SD cats at the same humane endpoint (n≥3).

A 10-point scale based on gait dysfunction in untreated SD cats (*8*) was used to track disease progression. Age of symptom onset ± standard deviation are shown in parentheses: 10, normal (<1.3 ± 0.2 mos.); 9, hind limb weakness (2.1 ± 0.0 mos.); 8, wide stance (2.2 ± 0.4 mos.); 7, ataxia (2.5 ± 0.3 mos.); 6, instability with occasional falling (2.9 ± 0.5 mos.); 5, limited walking (3.5 ± 0.5 mos.); 4, can stand but not walk (3.9 ± 0.5 mos.); 3, cannot stand (humane endpoint, 4.3 ± 0.2 months for untreated SD cats). One point was subtracted for acquisition of each of the following : fine tremors, full-body tremors, hind limb spasticity, fore limb spasticity.

### Tissue Preparation

Brains were divided into 9 transverse blocks of 0.6 cm from the frontal pole through the cerebellum (Figure 5). Blocks from the right hemisphere were frozen in optimal cutting temperature (OCT) medium and used for measuring specific enzyme activity by synthetic fluorogenic substrates based on 4-methylumbelliferone (4MU) as well as qPCR for AAV vector biodistribution. Blocks from the left hemisphere were halved to 0.3 cm and either flash frozen in liquid nitrogen or fixed in 10% formalin. Frozen tissue was stored at -80°C and used for high-performance thin layer chromatography (HPTLC) assays to quantify storage material. Formalin-fixed tissue was embedded in paraffin blocks and used for LFB (Luxol Fast Blue) or IHC staining.

### MRI and MRS

Cats were placed under general anesthesia with dexmedetomidine and ketamine and maintained with isoflurane. MRI and MRS data were collected as previously described (*37, 55*) on a 7 Tesla scanner (Siemens Healthcare) on SD cats immediately before necropsy (n=3 for untreated, n=15 for all treated cats) and on normal cats at 4-5 months, 8 months, and 12 months of age (n=3 per time point). To acquire MRS data, voxels were placed in the thalamus (7 × 6 × 8 mm), parietal cortex (7 × 6 × 8 mm), and cerebellum (7 × 7 × 8 mm). MRS data were processed with LC model and internal water scaling (http://www.s-provencher.com/lcmodel.shtml).

### Cerebrospinal Fluid (CSF) Analysis

Cats were placed under general anesthesia as described earlier for MRI and MRS acquisition. CSF was collected from the cisterna magna and immediately frozen at -80°C. A Cobas C311 chemistry analyzer (Roche Hitachi) was used to measure AST (aspartate aminotransferase) and LDH (lactate dehydrogenase). Acquisition of Hex specific activity is described below.

### Enzyme Activity Assays

Several frozen sections were cut from the OCT-frozen blocks and homogenized manually in 50 mM citrate phosphate buffer (pH 4.4) as previously described (*39*). The activity of HexA, Total Hex, Bgal, and Mann enzymes were measured with 4MU-based fluorogenic substrates as previously described (*39*), except 10µL was used for Bgal and Mann, 5µL for Total Hex, and 20µL for HexA. Specific activity was normalized to protein concentration through the Lowry method, then expressed as nmol 4MU/mg/hour.

### Lipid extraction and ganglioside quantification

Punch biopsies 8mm in diameter were taken from frozen CNS sections and lyophilized overnight. The HPTLC protocol was performed as previously described (*28*).

### Ultrasound elastography

Tissue elasticity was assessed with 2D shear wave elastography with acoustic radiation impulse force (ARIF) as previously described in the feline liver (*36, 56*), kidney (*35, 56*), and spleen (*56*). The semitendinosus was also assessed as a representative of skeletal muscle. Cats were sedated as needed while a Toshiba Aplio 500 ultrasound machine with a15L5 linear probe was applied to the organ of interest. Shear wave speed was quantified as Young’s modulus to determine tissue elasticity value (kPa). For the liver, measurements were recorded from 6-8 intralobular sites from the right, central, and left portions, which were then averaged together. Large vascular and ductal structures were avoided. Care was taken so that shear wave sampling ROI and depth were consistent between cats.

### Luxol Fast Blue Staining

Slides underwent a rehydration series of 8 minutes in Hemo-D (Scientific Safety Solvents, Keller, TX, USA) or xylene (3x), 2 minutes in 100% ethanol (2x), 2 minutes in 95% ethanol (2x), 2 minutes in distilled water, then incubated overnight at room temperature in LFB solution (0.1% in 95% alcohol with acetic acid, Electron Microscopy Sciences, Hatfield, PA, USA). In groups of 3 or 4 with at least one normal sample, slides were dipped in lithium carbonate (0.05%, Diagnostic BioSystems, Pleasanton, CA, USA) and 70% ethanol until the gray matter was clear. After a rinse in distilled water, slides were counterstained with Mayer’s Hematoxylin (Electron Microscopy Sciences) for 3 minutes, followed by a 3-minute rinse in running water. Slides dried overnight at room temperature and were coverslipped the following day.

### Immunohistochemistry

Slides were dehydrated in the same series as for LFB staining, then underwent antigen retrieval with Antigen Decloaker (Biocare Medical, Concord, CA, USA) in a decloaking chamber (Biocare Medical). After rinsing in distilled water, slides were loaded into an IntelliPath FLX slide stainer (Biocare Medical). The detailed protocol can be found in Supplemental Materials and Methods. All reagents were manufactured by Biocare Medical unless otherwise noted.

### Image Acquisition of Stained Slides

A Keyence BZ-X810 (Keyence, Itasca, Illinois, USA) was used to acquire all images. After brightness correction and scout scans at 4x, relevant regions were outlined at 20x and individual images were acquired. Images were stitched together with the built-in software and saved in the Big TIFF format. Files were imported into QuPath software (*57*) version 0.2.3 where they were qualitatively assessed, with representative images exported for inclusion in figures.

### Quantification of Stained Slides

After images were acquired and imported into the QuPath software (described above), algorithms were designed for each stain and tested on 2-3 images from cats in different treatment groups. For the nuclear stain Olig2, the algorithm was designed to count the number of stained nuclei per mm^2^ by using the cell detection function based on optical density, then applying an object classification to detected cells to differentiate DAB-positive cells from hematoxylin-positive cells if necessary. For the cytoplasmic IHC stains (GFAP, Iba-1) and LFB, algorithms were simple thresholders to calculate the percentage of stained pixels. See Supplementary Materials for algorithm design details.

### Histopathology

Formalin-fixed paraffin embedded tissue (described above) was stained with Hematoxylin & Eosin and evaluated by a board-certified veterinary anatomic pathologist who was blinded to treatment groups (AAV-treated n=15, untreated SD n=1, normal n=1).

### Quantitative PCR

Quantitative PCR was performed as previously described (*16*) on 50ng of genomic DNA using SsoAdvanced Universal SYBR Green Supermix (Bio-Rad) and QuantStudio 3 real-time PCR system (ThermoFisher). Primers were used from the SV40 polyadenylation site, results were standardized to feline albumin.

### Serum antibody titers

Serum antibody titers were evaluated as previously described (*16, 51*).

### Data Analysis and Statistics

All statistical analyses and graphs were generated with GraphPad Prism version 9.1.2 for Mac OSX (GraphPad Software, La Jolla California USA, www.graphpad.com). All tests were upaired T-tests with alpha values of 0.05.

## Author contributions

Conceptualization: DRM, MSE, TNS, HLG

Methodology: ASM, LT, DEO, ALG, JSC, PIH

Investigation: ASM, MS, ALG, HLG

Funding acquisition: MSE, DRM

Project administration: ASM, DRM, MSE, TNS

Supervision: ASM, DRM, ALG, TNS, MSE, HLG

Writing – original draft: ASM, DRM

Writing – review & editing: All

## Data and materials availability

All data are available in the main text or the supplementary materials.

## Supplementary Materials and Methods

### Immunohistochemistry detailed protocol

- General protocol:

1. Block: IP FLX Peroxidase, 5 minutes
2. Block: IP FLX Background Punisher, 5 minutes
3. Primary antibody: Diluted in Da Vinci Green Diluent
4. Secondary antibody: MACH 2 Rabbit HRP-Polymer, 30 minutes
5. Chromogen: Betazoid DAB, 5 minutes
6. Counter-stain: Tacha’s Auto Hematoxylin, 3 minutes
- Buffer wash throughout: TBS Auto Wash Buffer
- GFAP primary antibody: 1:200 dilution, 20 minutes
- Iba-1 primary antibody: 1:750 dilution, 30 minutes
- Olig2 primary antibody: Abcam, 1:200 dilution, 30 minutes Slides dried overnight at room temperature and were coverslipped the following day.

### QuPath workflow

The first step in the analysis of each image was to manually input the magnification as 20x and pixel length as 0.2 × 0.2 µm, since that information was not imported automatically with the metadata attached to each slide. Second, a stain vector was applied, if needed, to optimize the software’s differentiation between hematoxylin and DAB or LFB (stain vectors provided below). Third, regions of interest (parietal cortex white matter/gray matter, thalamus, cerebellar cortex white matter/gray matter, DCN, brainstem) were hand-drawn using the brush and wand tools. For LFB-stained slides, determination of white/gray matter boundaries during this step was performed while viewing the hematoxylin-only “channel” to avoid bias introduced by LFB staining. Fourth, the algorithm was applied and the resulting data manually recorded in a Microsoft Excel file.

### QuPath algorithms

- Olig2

○ Stain vectors:

▪ Hematoxylin: 0.739 0.538 0.405
▪ DAB: 0.452 0.589 0.67
▪ Background: 240 240 240
○ Cell detection:

▪ Detection image: Optical density sum
▪ Requested pixel size: 0.2 µm
▪ Background radius: 10 µm
▪ Median filter radius: 0 µm
▪ Sigma: 1.0 µm
▪ Minimum area: 6 µm^2^
▪ Maximum area: 400 µm^2^
▪ Threshold: 0.25
▪ Maximum background intensity: 2.0
▪ Split by shape: true
▪ Exclude DAB: false
▪ Cell expansion parameters: 0 µm
▪ Include cell nucleus: true
▪ Smooth boundaries: true
▪ Make measurements: true
○ Object classification

▪ Channel filter: DAB
▪ Measurement: DAB OD Mean
▪ Threshold: 0.08
▪ Above: Positive
▪ Below: Negative
- NeuN

○ Stain vector (faint hematoxylin)

▪ Hematoxylin: 0.50236 0.60145 0.6212
▪ DAB: 0.31918 0.51223 0.79734
▪ Background: 223 223 220
○ Stain vector (strong hematoxylin)

▪ Hematoxylin: 0.607 0.628 0.488
▪ DAB: 0.401 0.573 0.715
▪ Background: 221 223 221
○ Cell detection (faint hematoxylin)

▪ Detection image: Optical density sum
▪ Requested pixel size: 0.2 µm
▪ Background radius: 18 µm
▪ Median filter radius: 0 µm
▪ Sigma: 1.5 µm
▪ Minimum area: 6 µm^2^
▪ Maximum area: 600 µm^2^
▪ Threshold: 0.3
▪ Maximum background intensity: 2.0
▪ Split by shape: false
▪ Exclude DAB: false
▪ Cell expansion parameters: 3 µm
▪ Include cell nucleus: true
▪ Smooth boundaries: true
▪ Make measurements: true
○ Cell detection (strong hematoxylin)

▪ Same as faint hematoxylin except threshold = 0.4
○ Object classification (faint hematoxylin) #1

▪ Object filter: Detections (all)
▪ Channel filter: Hematoxylin
▪ Measurement: Nucleus Hematoxylin OD Mean
▪ Above: Negative
▪ Below: Positive
○ Object classification (faint hematoxylin) #2

▪ Object filter: Detections (all)
▪ Channel filter: DAB
▪ Measurement: Nucleus DAB OD Max
▪ Above: Positive
▪ Below: Negative
○ Object classification (strong hematoxylin)

▪ Object filter: Detections (all)
▪ Channel filter: DAB
▪ Measurement: Nucleus DAB OD Mean
▪ Threshold: 0.15
▪ Above: Positive
▪ Below: Negative
○ Stain vector (for strong-hematoxylin slides, cerebellar cortex only)

▪ Hematoxylin: 0.72 0.641 0.265
▪ DAB: 0.435 0.568 0.699
▪ Background: 217 220 220
- Calbindin

- Stain vector: Default
- Thresholder: same as NeuN thresholder, except Threshold = 0.30
- GFAP

- Stain vector: Default
- Thresholder (for strong-hematoxylin slides)

▪ Resolution: Very high
▪ Channel: DAB
▪ Prefilter: Gaussian
▪ Smoothing sigma: 0
▪ Threshold: 0.20
▪ Above threshold: Positive
▪ Below threshold: Negative
▪ Region: Any annotations
- Iba-1

○ Stain vector: Default
○ Thresholder: same as GFAP thresholder, except Threshold = 0.15
- LFB

○ Stain vector

▪ Hematoxylin: 0.644 0.722 0.251
▪ LFB: 0.777 0.516 0.359
▪ Background: 238 238 236
○ Thresholder:

▪ Channel: LFB
▪ Above threshold: Positive
▪ Below threshold: Negative

---

**Figure S1.**
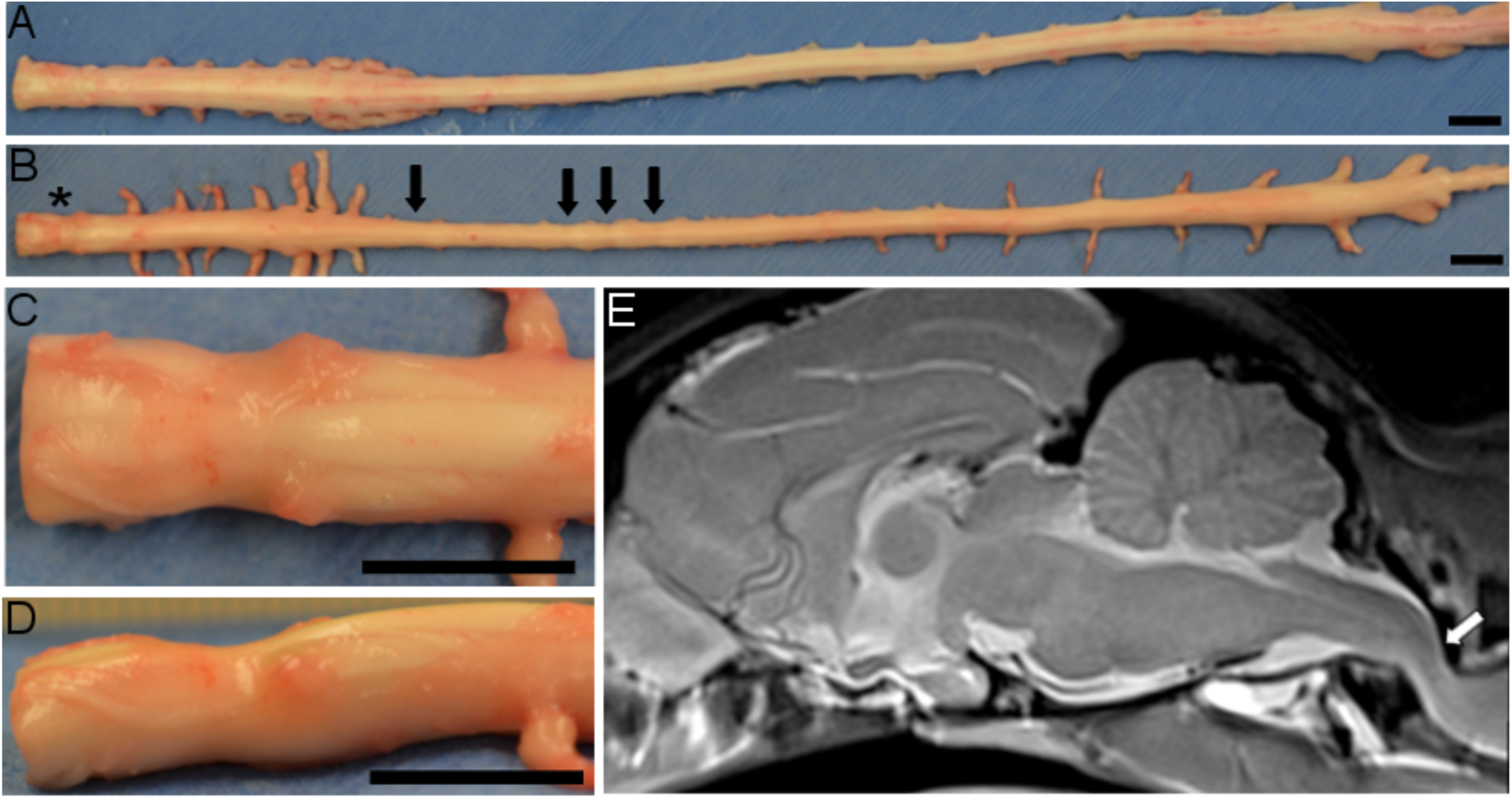
Spinal cord compression in AAV-treated cat. (A) Spinal cord of normal cat at necropsy. (B) Spinal cord of SD cat #52 of the high-dose cohort at necropsy (9.6 months old) with multiple compression lesions. Black arrows indicate sites of compression at T3 and T7-T9. * indicates most severe site of compression at C1, which is also depicted in (C-E). (C) Magnified view of dorsal surface of C1 compression lesion. (D) Magnified view of left lateral surface of C1 compression lesion. (E) Sagittal view of T2-weighted MRI on midline. White arrow indicates where the thin bright line of the CSF-filled central canal is interrupted at C1 due to spinal cord compression. Scale bar: 1cm.

**Figure S2.**
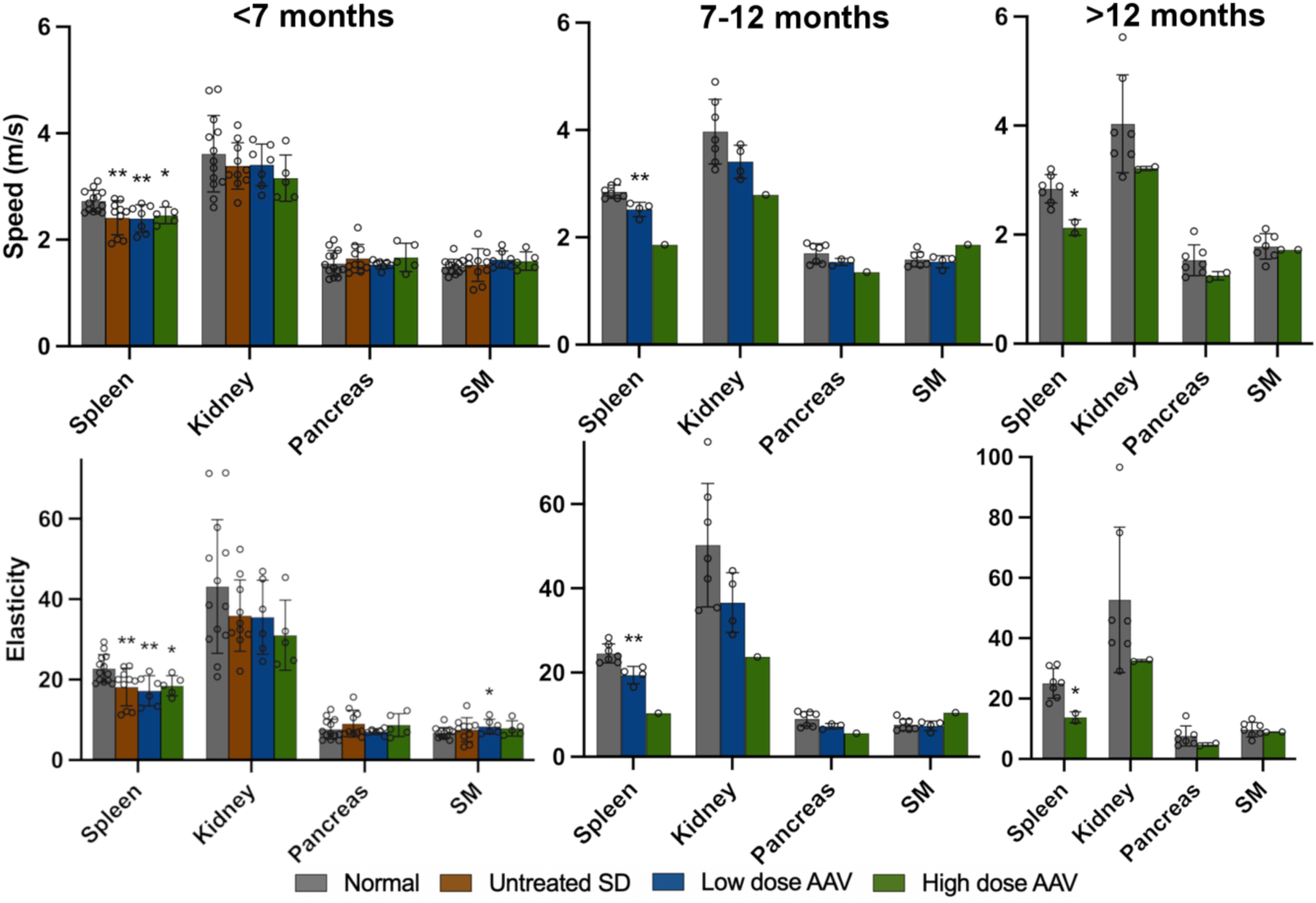
Ultrasound elastography. Shear wave ultrasound elastography results in the spleen, kidney, pancreas and skeletal muscle (SM). *p<0.05, **0.01>p>0.001 vs. age-matched normal cats.

**Fig S3.**
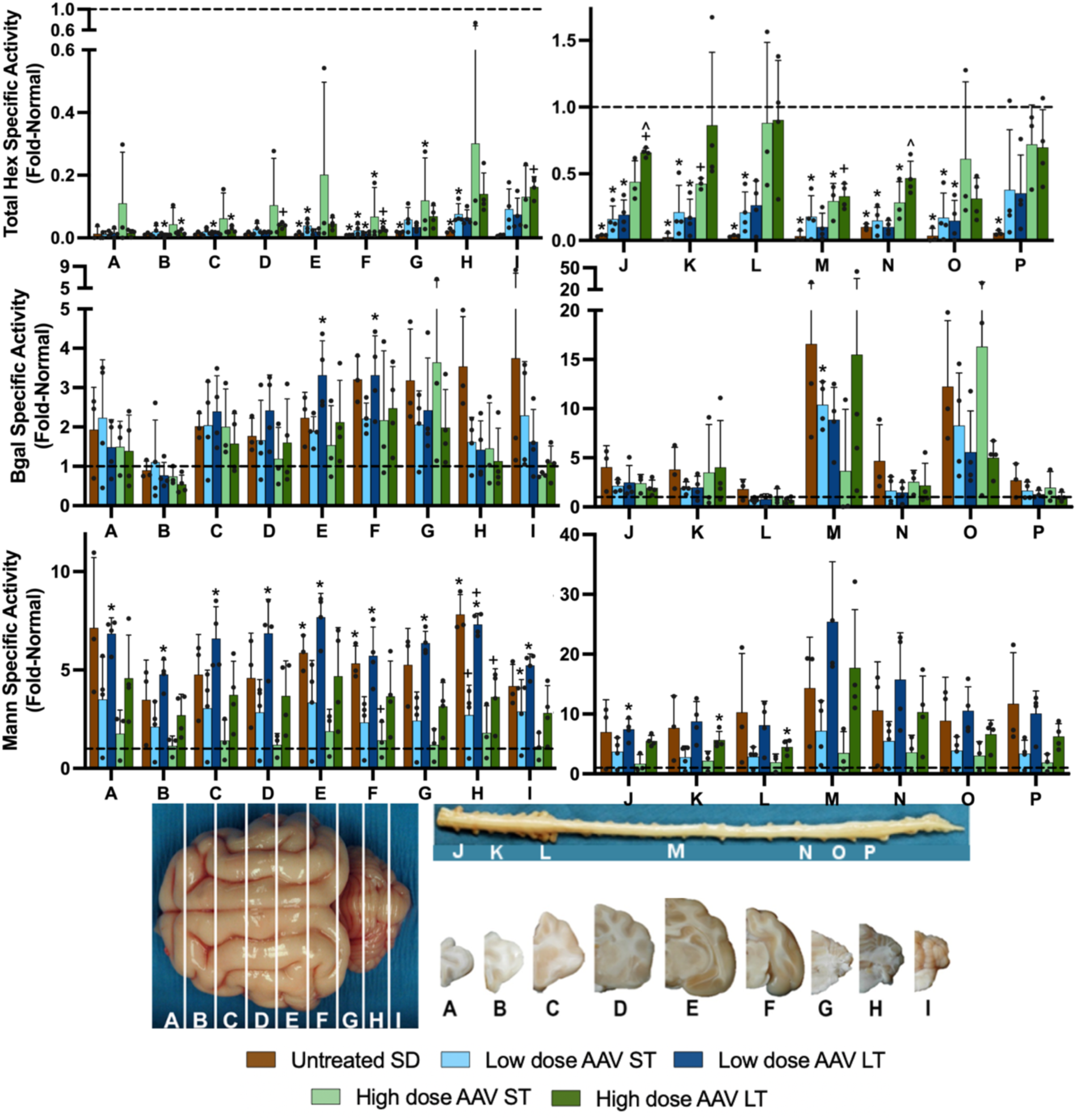
Specific activity of Total Hex, Bgal, and Mann lysosomal enzymes in the CNS. Specific activity of combined Hex isozymes (Total Hex) in the brain and spinal cord increases with AAV treatment in a dose-dependent fashion. Specific activity of other lysosomal hydrolases ß-galactosidase (Bgal) and α-mannosidase (Mann) increase to above-normal levels in SD cats in a compensatory attempt to reduce the abnormally high amount of storage material. Reduction of Bgal and Mann levels in AAV-treated cats corresponds with increases in HexT activity. * indicates p<0.05 compared to age-matched normal cats, + indicates p<0.05 compared to untreated SD cats, and ^ indicates p<0.05 compared to low dose at same age.

**Fig S4.**
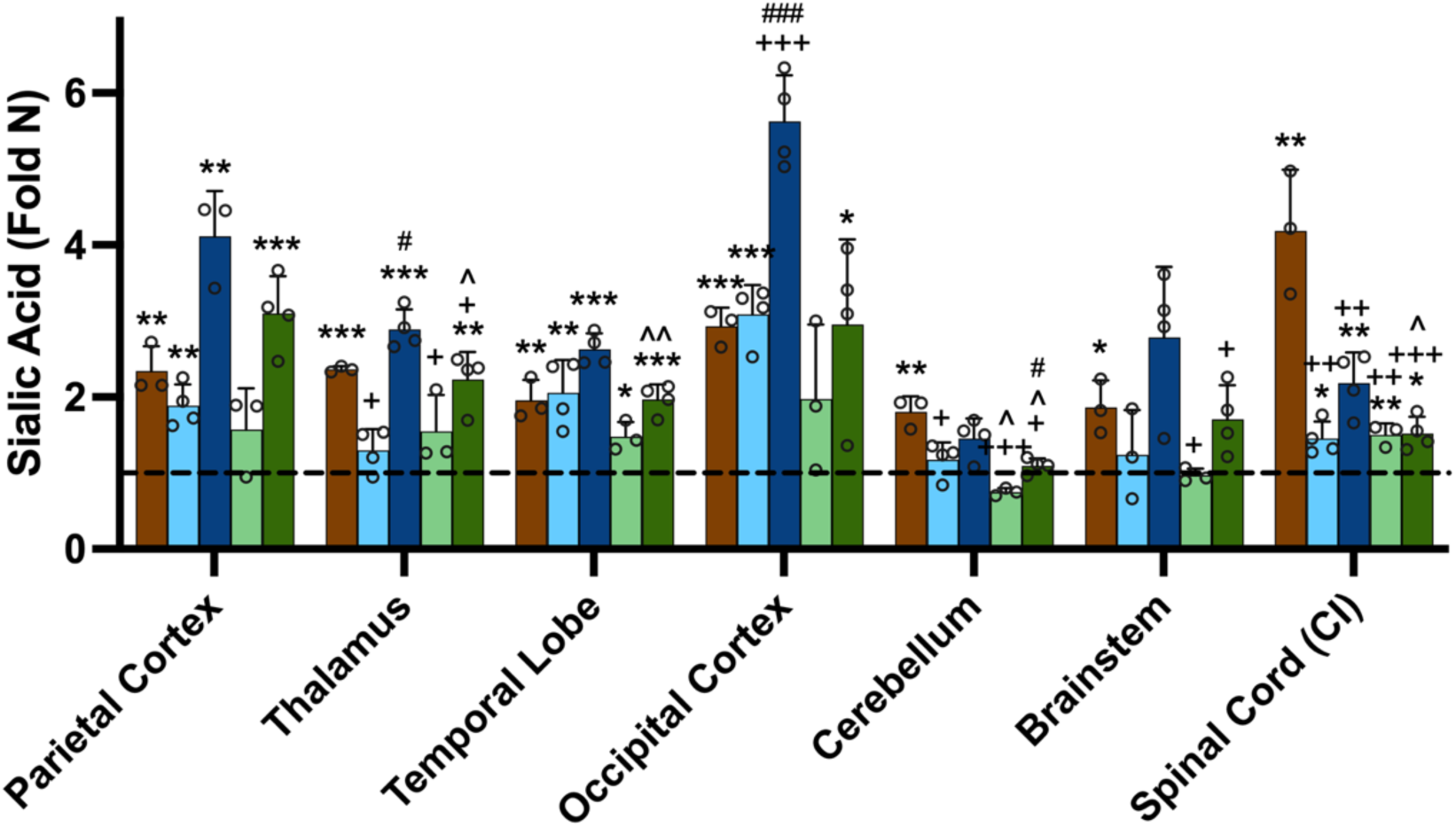
Total sialic acid storage products in AAV-treated cats. Total sialic acid (SA) is significantly increased above normal in 5/7 CNS regions examined. CI: cervical intumescence. *p<0.05, **0.01>p>0.001, ***0.001<p<0.0001 vs. age-matched normal cats; ^+^p<0.05, ^++^0.01>p>0.001, ^+++^0.001<p<0.0001 vs. untreated SD cats; ^0.05>p>0.01, ^^0.01>p>0.001 vs. low dose at same age. ^#^p<0.05, ^###^0.001<p<0.0001 vs. short term at same dose.

**Figure S5.**
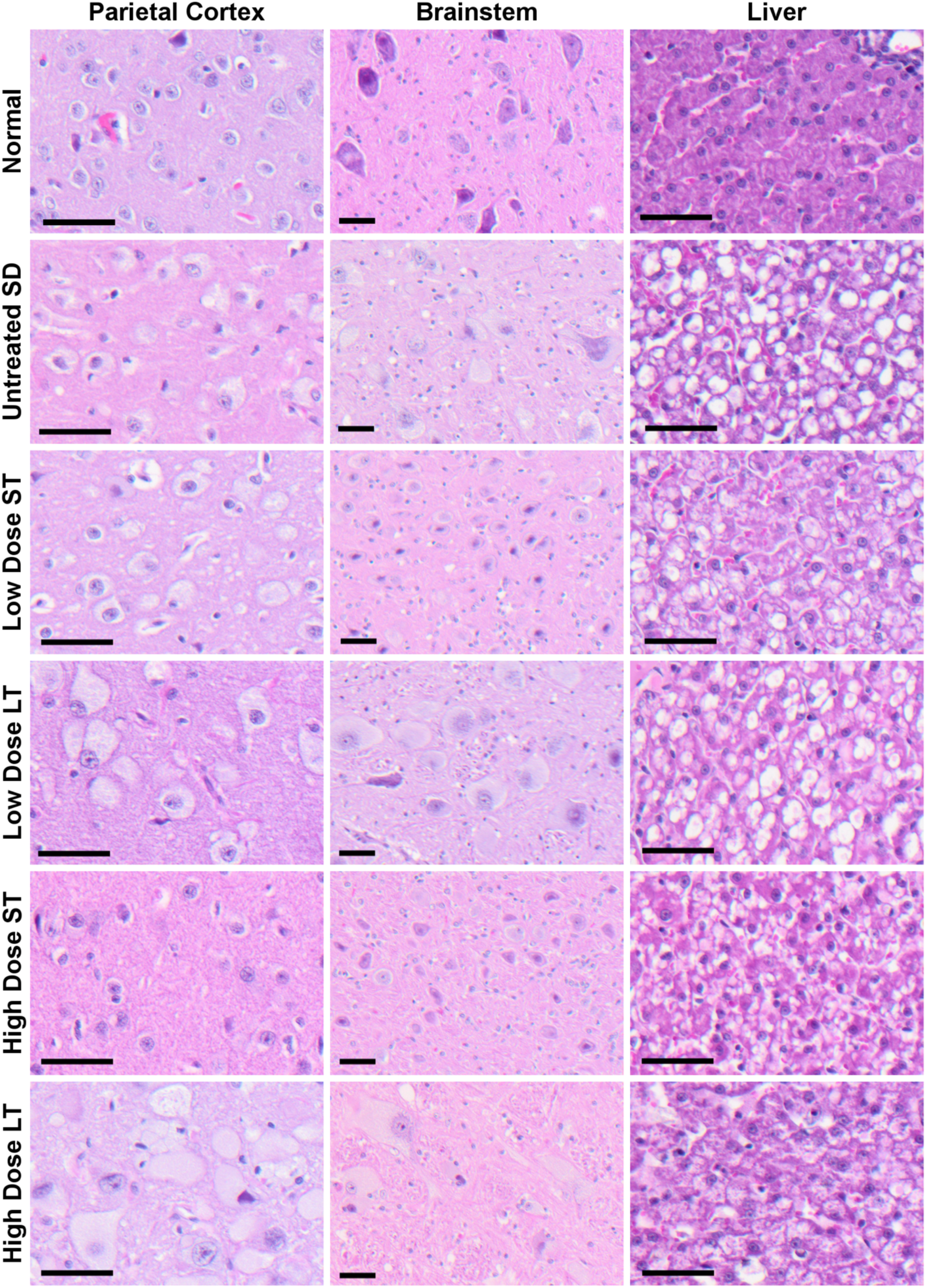
Histopathological storage lesions in neuronal cell bodies and hepatocytes are partially corrected by AAV treatment. Severe storage lesions occur frequently in the neuronal cell bodies and hepatocytes of untreated and AAV treated SD cats with a longer lifespan (long-term groups). Shorter-lived cats treated with the high dose demonstrated near-normalization of these lesions, with cats in the low dose group had moderate improvement. Micrographs at 10x (brainstem) or 20x (parietal cortex, liver) are from the representative cats. Scale bar: 25µm. ST: short-term, LT: long-term

**Fig S6.**
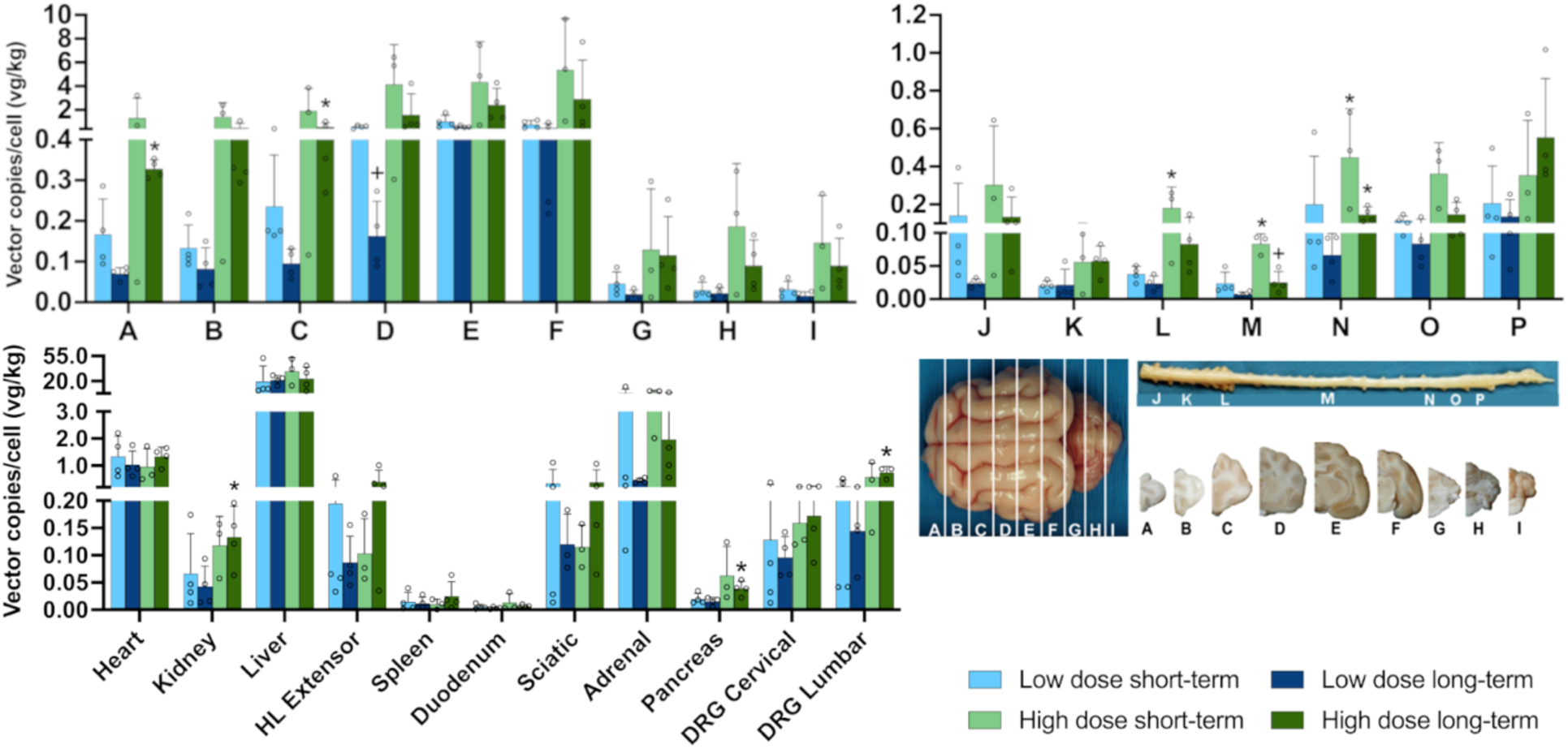
AAV vector biodistribution. AAV vector was distributed throughout the CNS and peripheral tissues, with the highest concentrations in the cerebral cortex and liver. ST: short-term, LT: long-term, HL: hindlimb, DRG: dorsal root ganglion. * indicates p<0.05 compared to age-matched low dose cats, + indicates p<0.05 compared to short-term cohort given the same dose.

**Fig S7.**
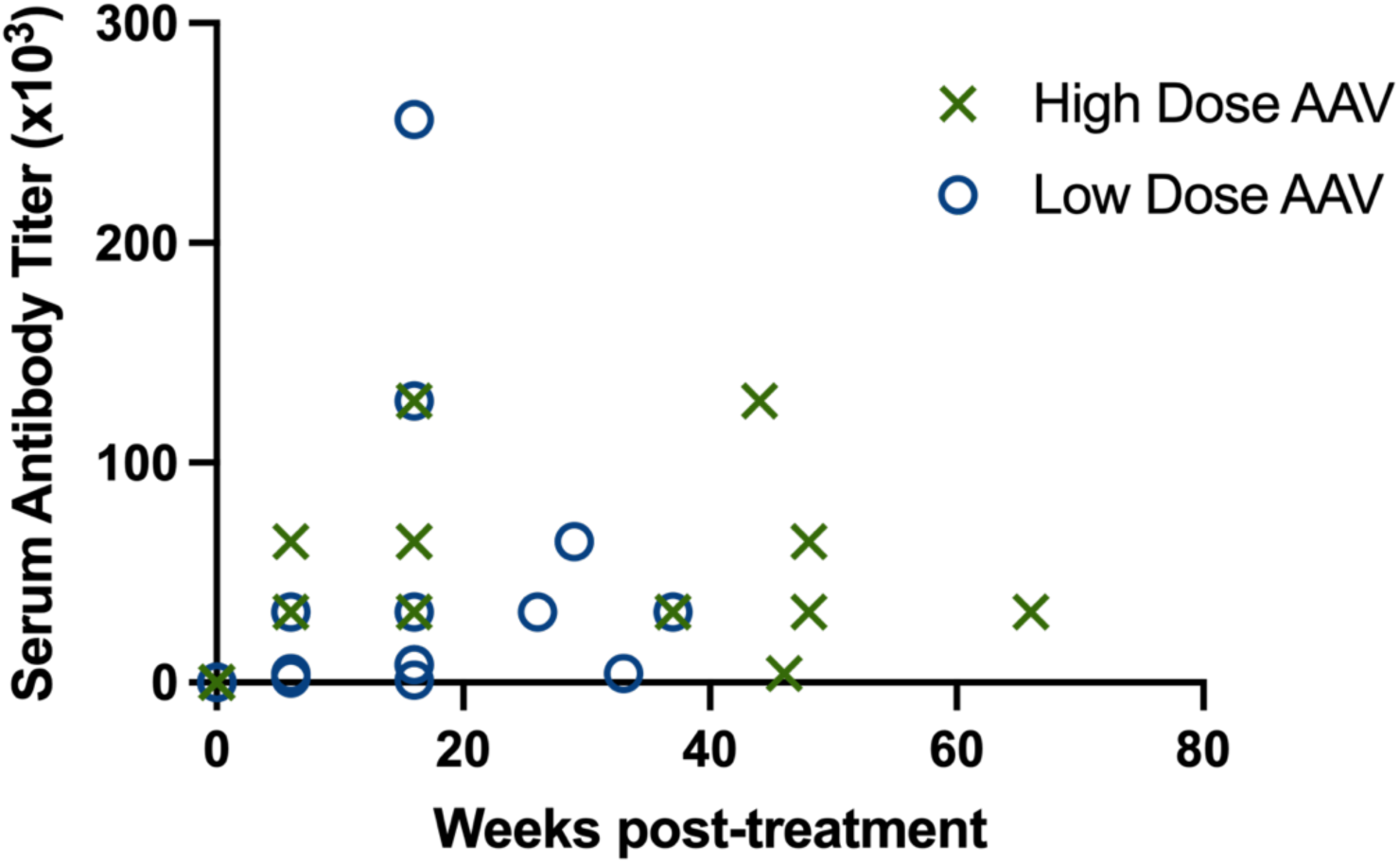
Serum antibody titers in AAV-treated cats. All cats developed antibody titers against AAV9. Overlapping data points within the same treatment group occur at 16 weeks (2 low dose cats with 8000 and 2 high dose cats with 64,000) and are not able to be distinguished on the graph.

